# The superior temporal asymmetrical pit remains asymmetric in autism spectrum disorder

**DOI:** 10.64898/2026.10.01.755796

**Authors:** Jake W. Wirtjes, Ethan H. Willbrand, Kevin S. Weiner

## Abstract

Neuroimaging studies identify the superior temporal sulcus (STS) as an anatomical and functional target in autism spectrum disorder (ASD). Yet, the STS is a heterogeneous structure containing unique subregions, such as the superior temporal asymmetrical pit (STAP), which is a recently identified human-specific landmark. To date, no study has compared the STAP between ASD individuals and neurotypical controls (NTs) in a large sample. Here, we analyzed the mean intra-hemispheric sulcal depth and inter-hemispheric laterality of the STAP, as well as the anterior and posterior portions of the STS (aSTS, pSTS), in 100 NTs and 100 ASD male individuals (ages 5-18) using a discovery-replication design. Across samples, the STAP and pSTS showed rightward asymmetry, while the aSTS showed leftward asymmetry within groups. All features of the STAP, aSTS, or pSTS did not differ significantly between groups. However, pSTS laterality related to individual differences in social cognition, which was further supported by a meta-analysis across over 200 studies showing that neurocognitive functional differences associated with social cognition and ASD are restricted to portions of the STS outside of the STAP. Together, these findings provide fine-grained insight into prior studies showing coarse morphological differences of the STS between individuals with ASD and NTs, as well as open up new questions and new neuroanatomical targets of the heterogeneous STS in future studies of brain structure, brain function, and cognition in ASD, as well as other neurodevelopmental disorders targeting the superior temporal cortex.

**Lay Summary:** Here, we show that a recently identified human-specific landmark within the superior temporal sulcus, a common target in ASD research, is comparable between ASD individuals and neurotypical controls. Yet, laterality metrics outside of this human-specific landmark correlate with individual differences in social cognition, which aligns with meta-analyses across hundreds of studies. These findings provide new neuroanatomical targets within the heterogeneous superior temporal cortex for future studies not only in ASD, but also other neurodevelopmental disorders.

## Introduction

Association cortices have expanded the most throughout evolution and contain structures that are human-specific, many of which have been linked to different neurodevelopmental disorders such as autism spectrum disorder (ASD; Ammons et al., 2021; Auzias et al., 2014; Brun et al., 2016; Cachia, A. et al., 2021; Fornito et al., 2006; Garrison et al., 2015; Nakamura et al., 2020; Parker et al., 2023; Ramos Benitez et al., 2024; Watanabe et al., 2014; <u>Willbrand et al., 2026a, 2026d</u>; Wu et al., 2026; Yucel et al., 2002). A common neuroanatomical target in ASD studies is the superior temporal sulcus (STS) within association cortex situated in the third visual, or lateral pathway. The STS is involved in a plethora of functions—from face, object, language, and number processing, to reading, multisensory integration, voice recognition, biological motion, and social cognition (Allison et al., 2000; Beauchamp et al., 2004; Belin et al., 2000; Binder et al., 1997; Brass et al., 2007; Calvert et al., 2001; Corbetta & Shulman, 2002; Deen et al., 2015; Fedorenko et al., 2012; Fletcher et al., 1995; Gallagher et al., 2000; Grossman et al., 2000; Haxby et al., 2000; Pelphrey et al., 2005; <u>Pitcher and Ungerleider, 2020</u>; Puce, 2024; Saxe & Kanwisher, 2003; Taylor et al., 2006; Vander Wyk et al., 2009; Vigneau et al., 2006). Previous studies identify abnormalities in the STS in ASD individuals, with consistent deficits in gray matter concentration, resting hypoperfusion, and atypical functional activity during social tasks (Boddaert et al., 2004; Ecker et al., 2013; Fan et al., 2025; Feng et al., 2025; Gervais et al., 2004; Ohnishi et al., 2000; Zilbovicius et al., 2000, 2006). Nevertheless, the STS is a large, functional and anatomical heterogeneous structure. For example, over a decade ago, Leroy and colleagues (2015) identified a human-specific landmark in the STS, the superior temporal asymmetrical pit (STAP), that is deeper in the right compared to the left hemisphere and has been linked to functional regions (Bodin et al., 2018). Yet, no study has compared the STAP in a large group of individuals with ASD relative to neurotypical controls (NTs), or examined the relationship between STAP morphology and laterality relative to individual differences in cognition relative to surrounding portions of the STS.

Zooming out to cortical morphology more broadly, deep sulcal points such as the STAP form early in gestation (Lohmann et al., 2008; Meng, Y. et al., 2014), and thus, have been hypothesized to act as a cerebral scaffolding for areas and networks, with developmental, clinical, and evolutionary implications (Cachia, A. et al., 2021; Natu et al., 2021; Rakic, 1988; Régis et al., 2005; Van Essen, 1997; <u>Willbrand et al., 2026b</u>; Zilles et al., 2013). As such, considering the STAP offers a first step in understanding the heterogeneity and “chaos” of the STS in ASD (Le Guen et al., 2018), as well as could further motivate future studies examining the relationship among sulcal morphology, functional networks, white matter architecture, and individual differences in cognition in ASD and other neurodevelopmental populations.

In their seminal study, Leroy and colleagues (2015) showed that there was no asymmetry in individuals with autism spectrum disorder (ASD). Nevertheless, the sample size was small (N=15 per group), necessitating a study with a larger group of participants that has yet to be conducted. Here, in 400 hemispheres (100 NT and 100 ASD individuals split evenly across discovery and replication samples), we found that the STAP was asymmetrical between the right and left hemispheres within and between groups. Intriguingly, the laterality of the posterior, but not the anterior, STS or the STAP related to individual differences in social cognition. Follow-up meta-analyses that included over 200 studies shows that neurocognitive functional differences associated with social cognition and ASD are restricted to portions of the STS outside of the STAP, further supporting the heterogeneity of the STS and the consistent morphology of the STAP between individuals with ASD and NTs. Together, these findings identify the STAP as a stable landmark in the superior temporal cortex that is similar between individuals with ASD and NTs, which highlights the importance of considering different portions of the STS when implementing morphological comparisons between different groups. Future studies should consider this human-specific landmark when analyzing data in the superior temporal cortex not only in ASD, but also in neurodevelopmental disorders more broadly.

## Methods

### Participants

Each included participant was randomly chosen from the Autism Brain Imaging Data Exchange (ABIDE; Di Martino et al., 2014). Because females are underrepresented in the ABIDE sample due to historical underdiagnosis, and given the sample’s emphasis on the developmental age range, participants were restricted to males under 20 years old. Following prior work (<u>Willbrand et al., 2026a</u>), we used a discovery–replication framework. The discovery sample comprised of 100 participants (50 NT, aged 10 ± 2.03 years; 50 ASD, aged 9.32 ± 2.73 years; p = .15), as did the replication sample (50 NT, aged 14.2 ± 3.05 years; 50 ASD, aged 13.1 ± 2.44 years; p = .053). All contributing sites confirmed IRB or ethics committee approval and HIPAA-compliant de-identification of shared data (Di Martino et al., 2014).

### Imaging Acquisition And Cortical Surface Reconstruction

All participants underwent a mock scanning session to become familiar with the scanning environment and process. A standard T1-weighted MPRAGE anatomical scan (TR = 2,530 ms, TE = 3.25 ms, 1.3 × 1 × 1.3 mm^3^ voxels) was acquired for each participant using a 3T Siemens Allegra MRI scanner at New York University, Georgetown University, and Stanford University (discovery sample), and at the University of Michigan (replication sample). Cortical surfaces were reconstructed using FreeSurfer (v6.0; Dale et al., 1999; Fischl et al., 1999). There was no effect of scanner site on sulcal morphology (all *p*s > 0.34).

Identification, Classification, and Feature Extraction of the Superior Temporal Asymmetrical Pit.

To localize the STAP in each hemisphere, an STS maximum probability map (MPM) was projected onto each participant’s native cortical surface (**Fig. 1A**, left). The STS MPM (Willbrand et al., 2024), defined in fsaverage space, was resampled to native space via FreeSurfer’s mri_label2label function and visually inspected and corrected as needed (**Fig. 1A**, middle). Importantly, this STS MPM was generated using previously published data that was independent from those examined in the present study. Sulcal depth was extracted from the FreeSurfer .sulc file (convexity; positive values denote sulcal vertices). The STAP was identified as the top 5% deepest vertices within the STS (**Fig. 1A**, right). See **Fig. 1B** and **Fig. 1C** for group average and example individual surfaces for NT and ASD participants, respectively. A laterality index (LI) was calculated to determine hemispheric asymmetry:

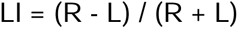

where L and R are STAP depth in the left and right hemispheres; more positive values indicate greater rightward asymmetry.

**Figure 1.**
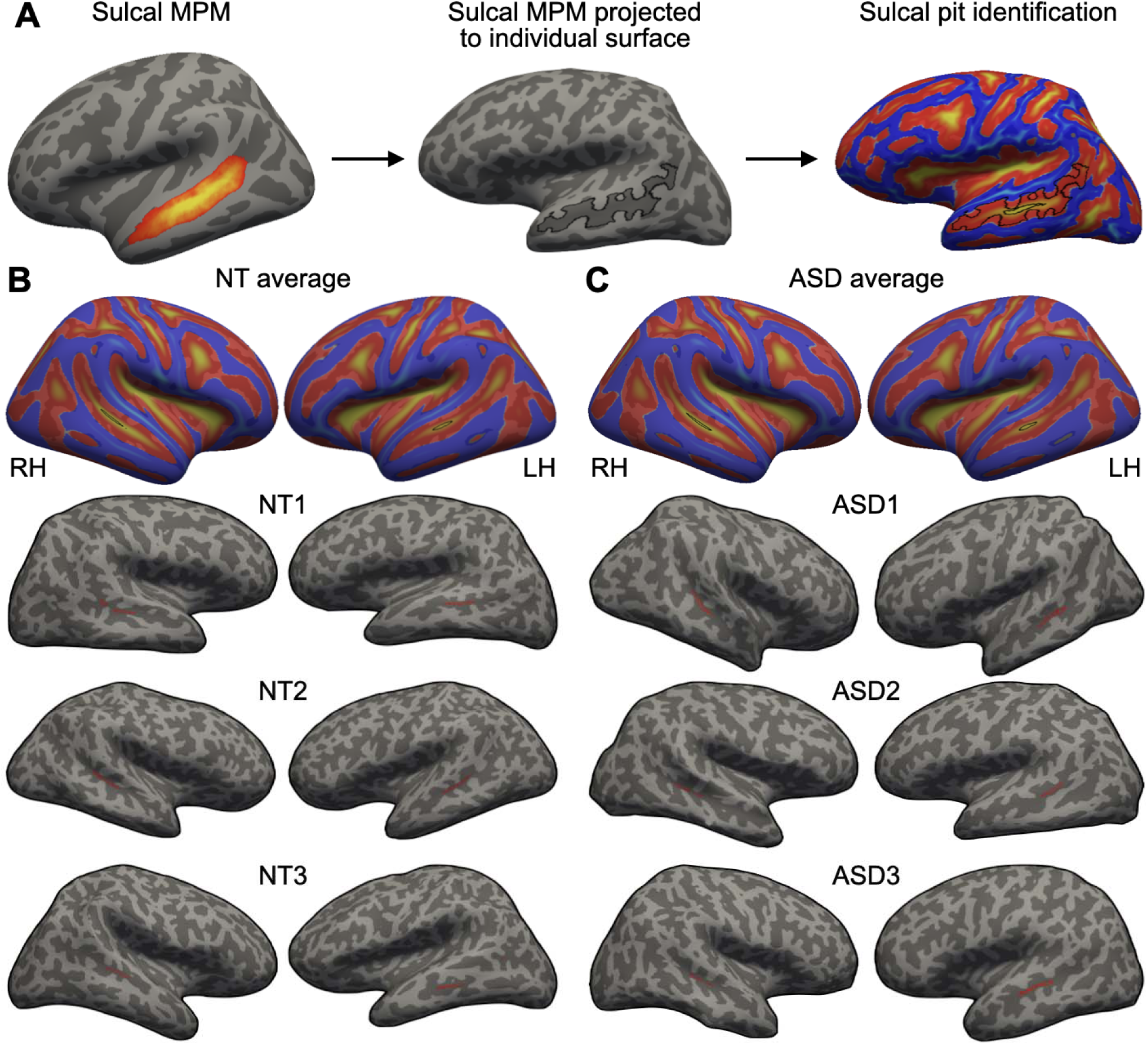
Identifying the superior temporal asymmetrical pit (STAP). **A.** Methodology to identify the STAP on the cortical surface (dark gray: sulci; light gray: gyri) in individual hemispheres. Maximum probability maps (MPM) of the STS (from Willbrand et al. (2024)) were projected to the native space of each participant and thresholded in each hemisphere (Methods for details). *Left:* STS MPM on the fsaverage surface. *Middle*: Projected STS MPM on a randomly selected individual left hemisphere. *Right*: The STAP (small black outline) identified as the top 5% of deepest vertices within the projected MPM STS label (large black outline). **B.** *Top row*: The average STAP (black outline) projected to left and right hemisphere inflated cortical surfaces relative to the unthresholded average sulcal map of the neurotypical (NT) participants included in the present study. *Bottom three rows*: three randomly chosen individual NT cortical surfaces with the STAP (red outline) overlaid. **C.** Same as **B.**, but for three randomly chosen individuals with autism spectrum disorder (ASD) used in the present sample.

### Segmentation of the STS into anterior and posterior portions

In each hemisphere, the STS was partitioned into anterior and posterior segments relative to each individual’s STAP (**Fig. 2A**). On the first principal component of the STAP (PC1; proxy for STS length), the most anterior and most posterior portions of the STAP were plotted, bounding the STAP. From this, all vertices anterior to the most anterior vertex of the STAP were labeled as the anterior portion of the STS. Similarly, all vertices posterior to the most posterior vertex of the STAP are labeled as the posterior portion of the STS. See **Fig. 2B** and **Fig. 2C** for example segmentation of the STS in NT and ASD individuals, respectively. Depth and laterality were extracted using the same process used for the STAP calculations.

**Figure 2.**
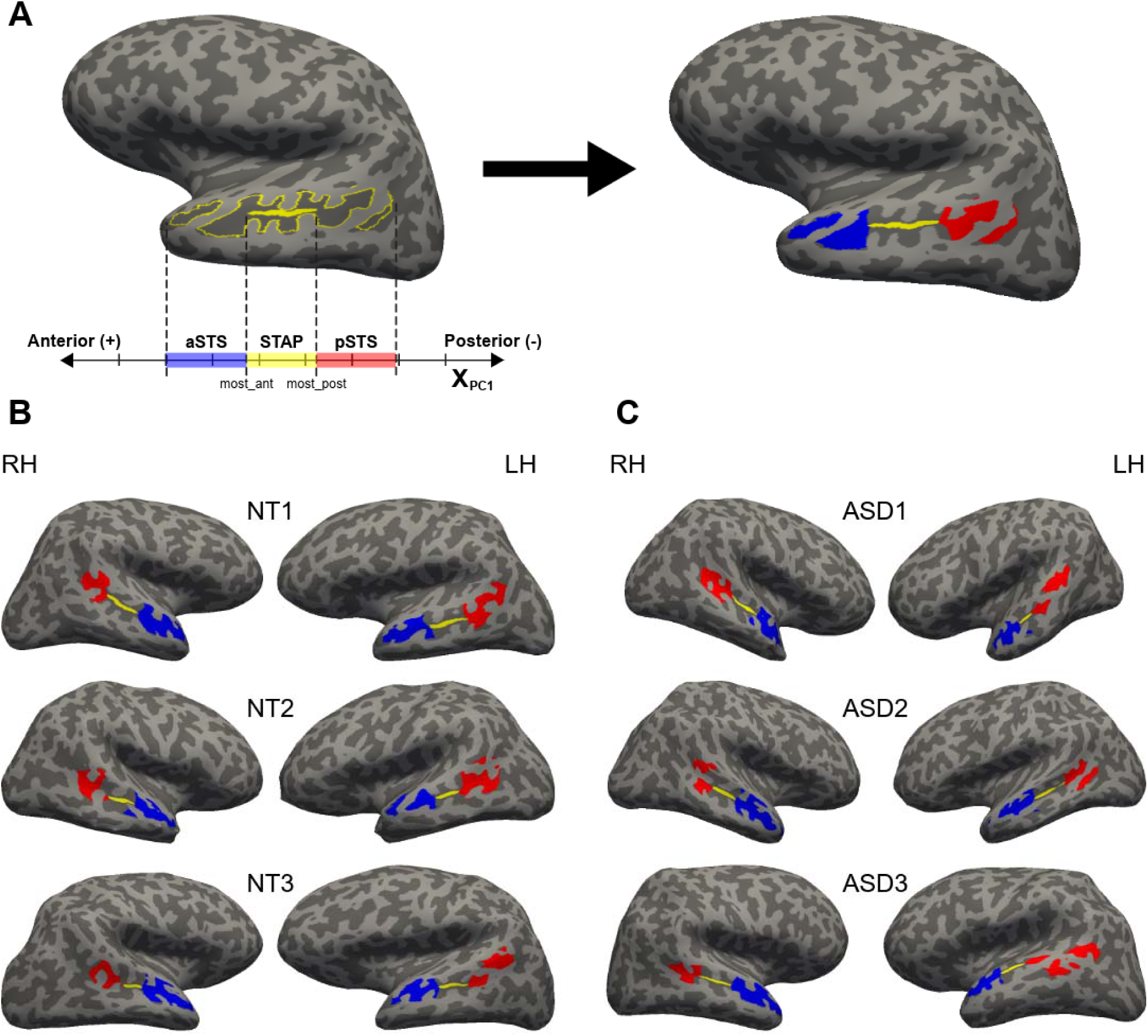
Methodology to segment anterior and posterior portions of the STS relative to the STAP in individual hemispheres. **A.** Example inflated left hemisphere detailing the segmentation of the anterior and posterior portions of the STS relative to the STAP at an individual level. *Left:* The STS (yellow; large, outline) and the STAP (yellow; small, filled) are plotted on the first principal projected axis (PC1; x-axis (black) indicating STS length). The most anterior (blue) and most posterior (red) portions of the STS label are identified on PC1 relative to the STAP (yellow; Methods for details). **B.** Three example individual NT cortical surfaces with the STAP (yellow), anterior (blue) and posterior (red) portions of the STS overlaid in the right (left column; RH) and left (right column; LH) hemispheres. **C.** Same as **B.**, but for individuals with ASD.

### Additional Sulcal Pits Examined

The following sulci were also examined beyond the STS: intraparietal sulcus (IPS; <u>Willbrand et al., 2026c</u>), inferior frontal sulcus (IFS; Willbrand et al., 2024), olfactory sulcus (OlfS; Hastings et al., 2024), parieto-occipital sulcus (POS; Maboudian et al., 2024), and transverse olfactory sulcus (tOlfS; Hastings et al., 2024). As for the STS, projected labels were restricted to sulcal vertices (curvature > 0). Pit identification and depth extraction followed the same procedure as the STAP (**Fig. 1A**). These sulci were selected as they lie within regions where previous studies suggested broad morphological differences in individuals with ASD (Auzias et al., 2014; Levitt et al., 2003; Nordahl et al., 2007; Ong & Fan, 2023; Tezcan et al., 2025), as well as functional regions and connectivity in these cortical expanses in individuals with ASD (Keown et al., 2013; Minshew & Keller, 2010; Stickel et al., 2019).

### Behavioral Metrics

Two standard ASD evaluation tests were considered: The Autism Diagnostic Observation Schedule (ADOS-2; Gotham et al., 2009; Lord et al., 2012) and the Social Responsiveness Scale (SRS, Constantino et al., 2003). Symptom severity metrics were available only in the discovery dataset; 45/50 and 46/50 ASD individuals had ADOS-2 and SRS scores reported, respectively. 47/50 NT individuals had SRS scores reported. NTs did not have ADOS-2 scores.

### Relating Neurosynth Meta-analysis Maps to the STS

To test potential relationships between the STS and the broad literature on ASD and ASD-related behaviors, we downloaded three association test maps (FDR-corrected χ*2* test, p = 0.01) from NeuroSynth (https://neurosynth.org/; Yarkoni et al., 2011): (i) *social cognition* (220 studies), (ii) *communication* (271 studies), and (iii) *autism* (244 studies). Each map was projected to *fsaverage* surface space with the *mri_vol2surf* FreeSurfer function to spatially relate them to the STS.

### Statistical Analyses

All analyses were conducted in the discovery and replication datasets separately. Three morphological features were analyzed (LI, left mean depth, and right mean depth). The proportion of individuals exhibiting rightward asymmetry (LI > 0) was compared between groups using Fisher’s exact test. Between-group differences (NT vs. ASD) were assessed using a Welch’s independent two-sample t-test. Within-group hemispheric differences were assessed using a paired-samples t-test. Group × region × hemisphere interactions were tested with an analysis of variance (ANOVA) model. Group differences were further tested with analysis of covariance (ANCOVA) models with group as the primary factor and age (both samples) and scan site (discovery sample) as covariates. Handedness was not considered due to incomplete recording. Outliers were removed within each distribution using an interquartile-range (IQR) filter (± 3×IQR; individuals below Q1 − 3×IQR or above Q3 + 3×IQR were excluded). Behavioral comparisons were corrected using the Benjamini-Hochberg false discovery rate (FDR) procedure.

## Results

### The superior temporal asymmetrical pit remains asymmetric in autism spectrum disorder

STAP asymmetry and mean STAP depth were similar between ASD and NT individuals (**Fig. 3 & S1**). STAP LI was compared between groups in the discovery and replication samples (**Fig. S1**). STAP mean depth showed a similar significant rightward lateralization observed in both groups (**Fig. 3A & 3B**; all p < .001), as well as no significant differences in corresponding hemispheres between groups (Discovery – Left: p = .58, Right: p = .69; Replication – Left: p = .86, Right: p = .20). Additionally, no significant differences in STAP LI (Discovery: p = .92; Replication: p = .49) or in the proportion of individuals exhibiting rightward asymmetry (Discovery: NT 78% vs. ASD 80%; Replication: NT 76% vs. ASD 78%; both Fisher’s exact p = 1.00) were found between groups (**Fig. S1**). Results remained consistent after adjusting for age and scan site (Discovery – LI: p = .89, Left: p = .95, Right: p = .84; Replication – LI: p = .24, Left: p = .57, Right: p = .40). Group × age interactions were not significant for all outcomes tested (all p > .32). Additionally, for both LI and sulcal depth (separately for each hemisphere), there was no relationship between morphological metrics and either ADOS-2 (LI p = .68, Left p = .20, Right p = .16; **Fig. S3 & S4**) or SRS (LI p = .16, Left p = .35, Right p = .09; **Fig. S3 & S4**) severity scores.

**Figure 3.**
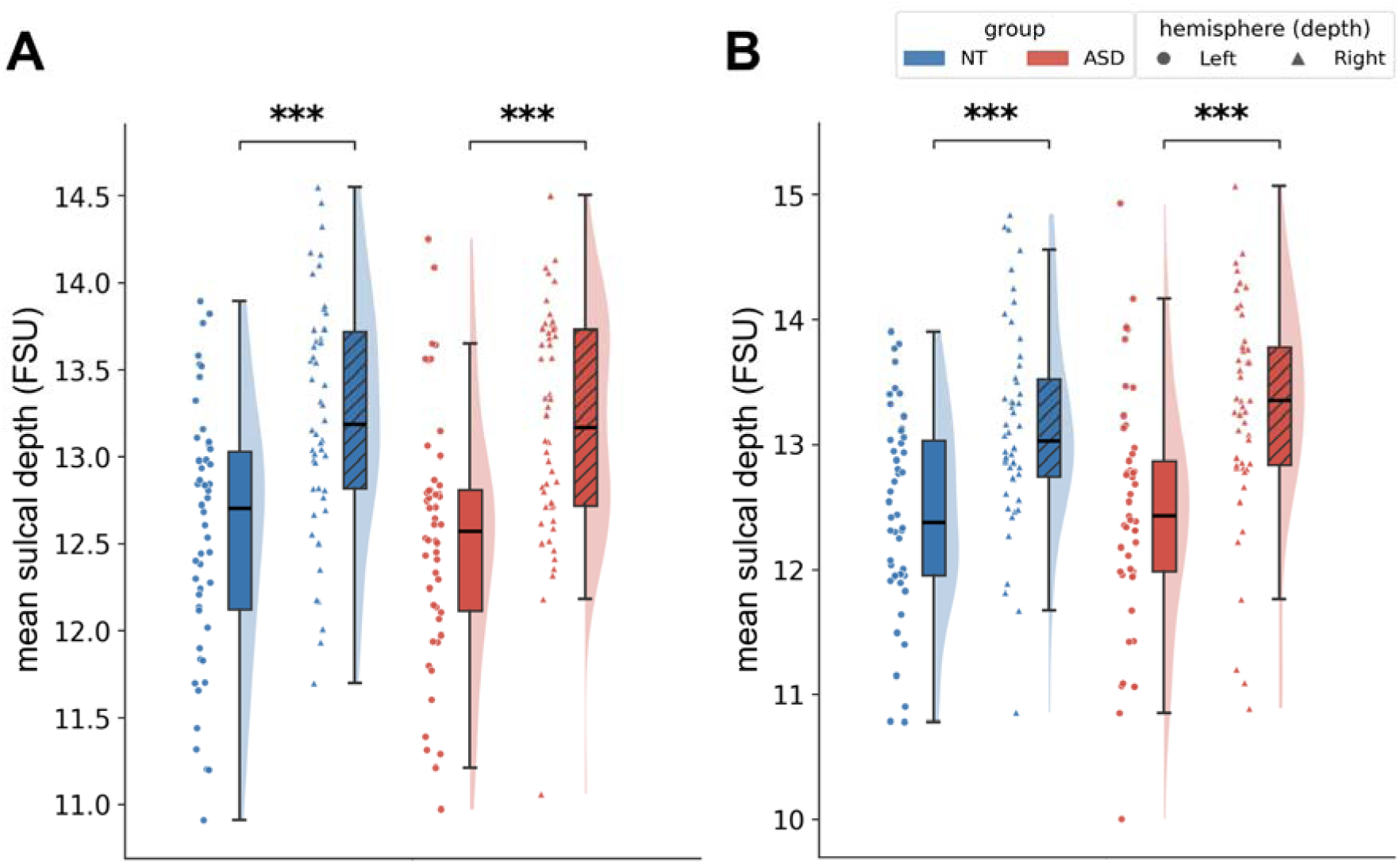
STAP morphology is conserved between ASD and NT individuals. **A**. Raincloud plots visualizing STAP mean depth in the left (circle) and right (triangle) hemispheres of NT (blue) and ASD (red) individuals. **B.** Same as **A.**, but in the independent replication dataset. In each sample (Discovery and Replication) and group (ASD, NT), the right hemisphere was significantly deeper than the left hemisphere (***p < .001) with no effect of group (all *p*s > .05). FSU = FreeSurfer units (Dale et al., 1999; Fischl et al., 1999).

### Left hemispheric bias in the anterior, but not posterior, STS outside the STAP

In addition to the STAP, we tested if the anterior and posterior portions of the STS outside the STAP were different between hemispheres and groups (**Fig. 4 & S1**). An ANOVA with region, hemisphere, and group as factors revealed a significant main effect of hemisphere (Discovery – F(1,96) = 12.32, p < .001; Replication – F(1,92) = 14.71, p < .001), as well as a significant region × hemisphere interaction (Discovery – F(1,96) = 34.98, p < 0.001; Replication – F(1,92) = 38.16, p < 0.001). Specifically, similar to the STAP, the posterior portion of the STS exhibits rightward asymmetry (Discovery – Left: 5.681 ± 2.401 FSU, Right: 7.105 ± 2.084 FSU, (R>L) p = 1.263 × 10^−5^; Replication – Left: 5.503 ± 2.156 FSU, Right: 6.84 ± 1.9 FSU, (R>L) p = 8.882 × 10^−6^; **Fig. 4 & S1**), while the anterior portion of the STS exhibits a leftward asymmetry (Discovery – Left: 4.704 ± 0.508 FSU, Right: 4.334 ± 0.483 FSU, (L>R) p = 1.662 × 10^−7^; Replication – Left: 4.636 ± 0.737 FSU, Right: 4.289 ± 0.64 FSU, (L>R) p = 1.559 × 10^−5^; **Fig. 4 & S1**). No differences between groups were found in laterality or depth in the discovery (group p = .29; group × hemisphere p = .50) or replication (group p = .60; group × hemisphere p = .80) samples (**Fig. 4 & S1**).

**Figure 4.**
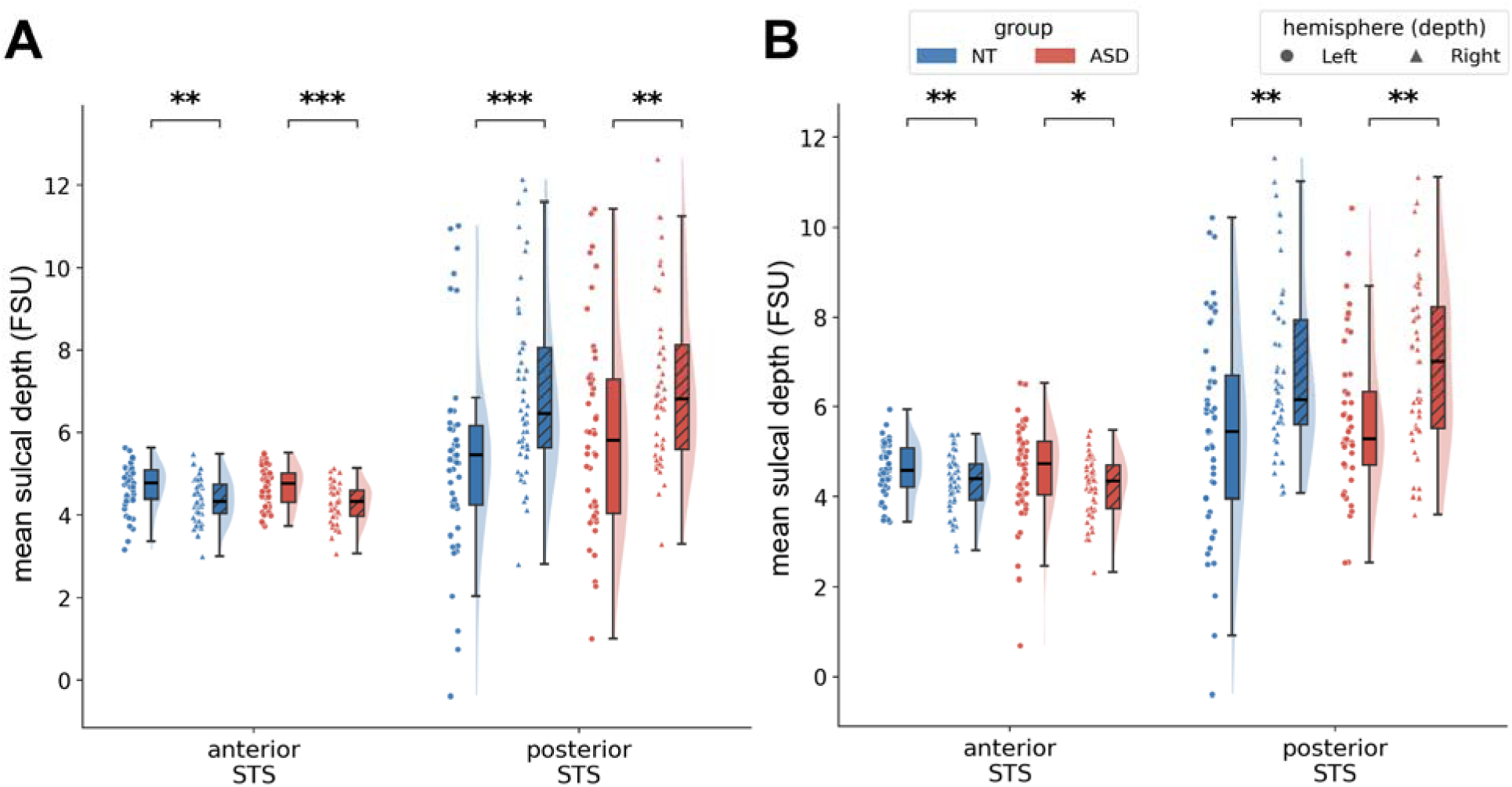
Outside the STAP: Left hemispheric bias in the anterior, but not posterior, STS. **A.** Raincloud plots visualizing mean depth of the anterior (left) and posterior (right) portions of the STS. Individual dots (left hemisphere) and triangles (right hemisphere) represent individual NT (blue) and ASD (red) participants. **B.** Same as **A.**, but in the independent replication dataset. In each sample (Discovery and Replication) and group (ASD, NT), there was a left hemispheric bias in the anterior, but not posterior, STS outside the STAP (***p < .001) with no effect of group (all *p*s > .05) and a hemisphere x region interaction (Discovery – F(1,96) = 34.98, p < 0.001; Replication – F(1,92) = 38.16, p < 0.001). Thus, the hemispheric asymmetry for the STAP and posterior STS is similar, while the bias in the anterior STS is in the opposite direction. FSU = FreeSurfer units (Dale et al., 1999; Fischl et al., 1999).

As previous findings showed relationships of functional regions and networks in olfactory, intraparietal, and occipital cortices in ASD (Keown et al., 2013; Minshew & Keller, 2010; Stickel et al., 2019), we tested if there were differences between metrics of sulci within these cortical expanses between groups. There were no significant differences between groups in either sample when considering FDR correction (**Fig. S2**). Specifically, there was no main effect of group for the pit of the IFS (Discovery – F(1,98) = 2.20, p = .14; Replication – F(1,98) = 0.35, p = .55), IPS (Discovery – F(1,98) = 3.54, p = .063; Replication – F(1,98) = 1.70, p = .20), OlfS (Discovery – F(1,98) = 2.67, p = .11; Replication – F(1,98) = 3.56, p = .062), or POS (Discovery – F(1,98) = 0.49, p = .49; Replication – F(1,98) = 0.011, p = .92). The tOlfs pit showed a significant main group effect with deeper pits in NT than ASD participants in the replication sample (NT – 4.03 ± 1.81 FSU; ASD – 3.07 ± 2.12 FSU; F(1,98) = 6.01, p = .016), but not in the discovery sample (NT – 3.28 ± 1.18 FSU; ASD: 2.78 ± 1.89 FSU; F(1,98) = 2.52, p = .12). There was no group × hemisphere interaction, indicating no group differences in pit asymmetry, for the IFS (Discovery – F(1,98) = 0.19, p = .66; Replication – F(1,98) = 0.003, p = .95), OlfS (Discovery – F(1,98) = 0.51, p = .48; Replication – F(1,98) = 0.80, p = .37), POS (Discovery – F(1,98) = 0.33, p = .57; Replication – F(1,98) = 3.86, p = .052), or tOlfS (Discovery – F(1,98) = 0.015, p = .90; Replication – F(1,98) = 1.72, p = .19).The IPS pit showed a significant group × hemisphere interaction in the replication sample (F(1,98) = 4.87, p = .030) but not in the discovery sample (F(1,98) = 1.41, p = .24).

Regarding hemispheric effects, the IFS pit was deeper in the left hemisphere (Discovery – F(1,98) = 15.49, p < .001; Replication – F(1,98) = 22.48, p < .001), whereas the pits of the OlfS (Discovery – F(1,98) = 8.86, p = .004; Replication – F(1,98) = 18.02, p < .001) and POS (Discovery – F(1,98) = 10.92, p = .001; Replication – F(1,98) = 10.42, p = .002) were deeper in the right hemisphere (**Fig. S2**). The IPS (Discovery – F(1,98) = 0.54, p = .46; Replication – F(1,98) = 1.52, p = .22) and tOlfS (Discovery – F(1,98) = 1.21, p = .27; Replication – F(1,98) = 2.34, p = .13) showed no hemispheric asymmetry (**Fig. S2**).

### Laterality of the posterior, but not anterior, STS correlates with SRS scores

Lastly, we compared ASD symptom severity scores (ADOS-2 and SRS) with sulcal features of the anterior and posterior portions of the STS (LI, left mean depth, and right mean depth) within the discovery dataset (Methods). We found that the laterality of the posterior portion of the STS is associated with SRS scores in ASD individuals (ρ = +.30, p = .049; **Fig. 5**). However, the laterality of the anterior STS was not correlated with SRS scores (p = .963; **Fig. 5**). Neither pSTS nor aSTS depth was correlated with SRS scores (Anterior: Left, p = .99; Right. p = .79; Posterior: Left, p = .23, Right, p = .12; **Fig. S6**). Furthermore, laterality and depth of the aSTS and pSTS in ASD individuals showed no correlation with ADOS-2 (Anterior – LI p = .14, Left p = .57, Right p = .11; Posterior – LI p = .11, Left p = .20, Right p = .37; **Fig. S5 & S6**). In the NT group, SRS scores were unrelated to laterality (all p > 0.26) or depth (all p > 0.08, all q > 0.16).

**Figure 5.**
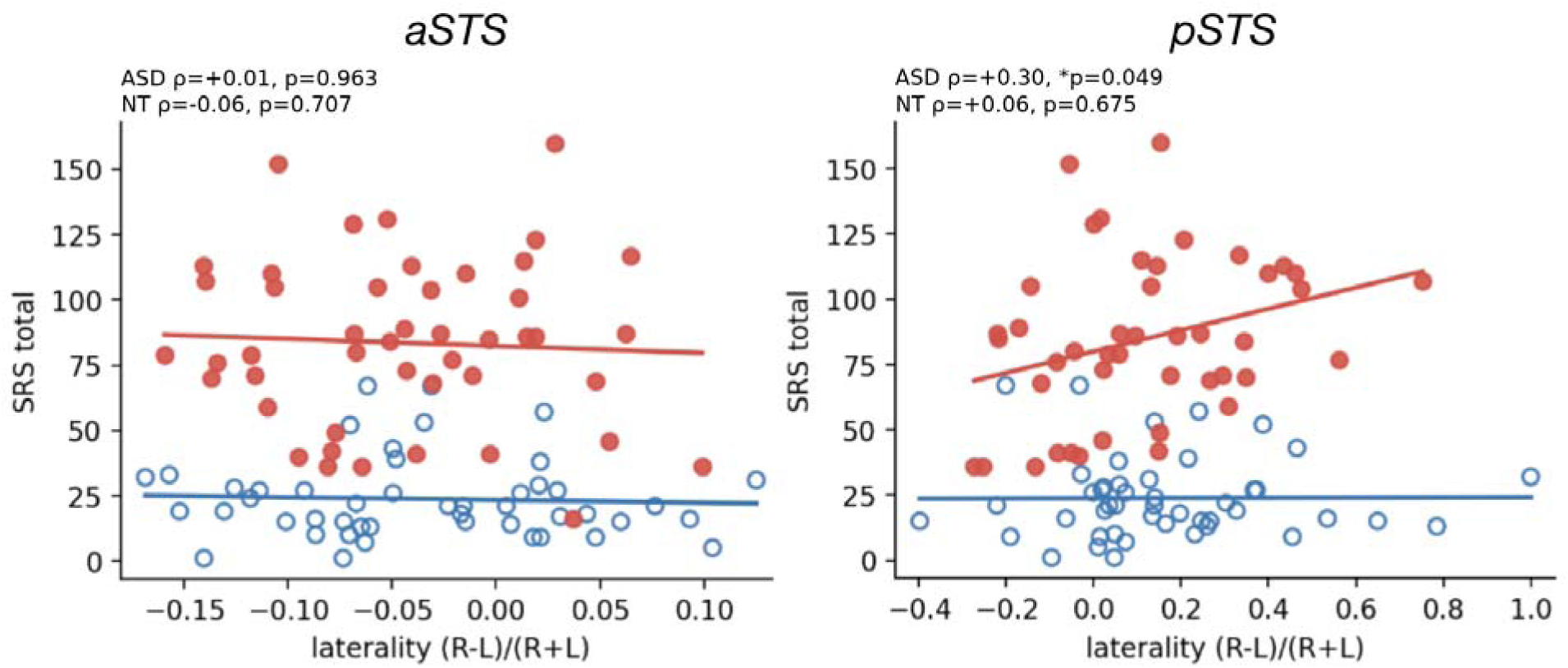
Laterality of the posterior, but not anterior, STS correlates with social cognition. Scatterplots visualizing the relationship between laterality (measured by the laterality index; LI) of the anterior (left) and posterior (right) STS and social cognition as measured by Social Responsiveness Scale (SRS) severity scores. Individual dots represent individual participants (ASD: red, filled; NT: blue, hollow). Solid red (ASD) and blue (NT) lines represent lines of best fit. Posterior STS laterality was found to correlate with SRS score in ASD individuals (ρ = +.30, *p = .049), while anterior laterality did not (p = .963). Anterior and posterior STS laterality in NTs did not show any association with SRS scores (all *p*s > .05). ρ = Spearman’s correlation coefficient. Significant *p < .05, Not significant p > .05

## Discussion

In the present study, we examined STAP morphology in young male ASD individuals and NTs. We quantified STAP morphology in two independent samples totalling 100 NT and 100 ASD participants. Across samples, the STAP and pSTS showed rightward asymmetry, while the aSTS showed leftward asymmetry within groups. All features of the STAP, aSTS, or pSTS did not differ significantly between groups. However, pSTS laterality related to individual differences in social cognition. We discuss these results in the following three sections: (i) STS, but not STAP, differences in ASD individuals and NTs, (ii) novel anatomical and functional targets in ASD research, and (iii) developmental trajectory of neuroanatomical structures as predictors for future studies, as well as limitations and future directions.

### STS, but not STAP, differences in ASD individuals and neurotypical controls

Given previous findings showing anatomical and functional differences in the STS between ASD individuals and NTs (Cauda et al., 2017; Guo et al., 2024; Keown et al., 2013; Minshew & Keller, 2010; Stickel et al., 2019; Zilbovicius et al., 2006), our findings may seem at odds with these prior studies. Instead, it is likely that the differences between groups are driven by portions of the STS outside of the STAP, for instance, in the posterior or anterior portions of the STS. Our analyses provide anatomical support for this hypothesis as the anterior and posterior STS showed different asymmetries in which the latter showed a rightward asymmetry, while the former showed a leftward asymmetry. Further, previous studies show that the STAP has been linked to underlying genetics, while other portions of the STS are likely under less genetic control (Le Guen et al., 2018, 2020), proposing that the STAP is more broadly involved in human communication, while posterior and anterior portions of the STS are involved in higher-level aspects of communication involving social interaction.

To explore this possibility, we performed a meta-analysis in NeuroSynth (Yarkoni et al., 2011). Using the search term “social cognition” across 220 studies resulted in several clusters along the STS (**Fig. 6**). These clusters were in the anterior and posterior portions of the STS, but not in the middle of the STS, where the STAP is located. Additionally, there could even be a more fine-grained functional parcellation as prior work differentiated the pSTS into three functional subregions in ASD individuals using a data-driven functional connectivity approach (Shih et al., 2011). As pSTS laterality correlated with SRS scores in the present study, it would be interesting for future studies to test if this relationship is driven by a more fine-grained parcellation of the pSTS. Further, a recent study showed functional connectivity differences between pSTS and mSTS regions in ASD individuals compared to NTs (Li et al., 2025), emphasizing that this cortical expanse is consistent with what has been referred to as the social visual pathway (Pitcher and Ungerleider, 2021).

**Figure 6.**
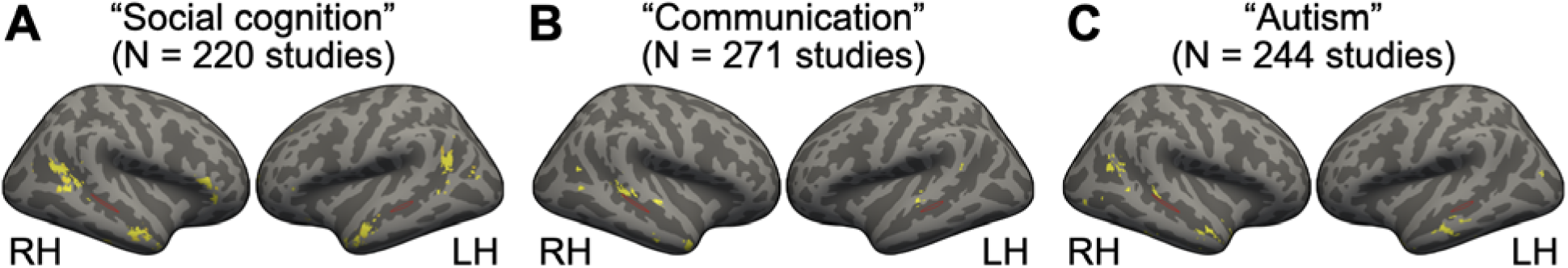
The STAP does not co-localize with Neurosynth meta-analytic clusters related to social cognition, communication, and autism terms. **A.** Left and right hemisphere inflated fsaverage cortical surfaces (left surface: right hemisphere, RH; right surface: left hemisphere, LH) displaying overlap of a whole-brain FDR-corrected (p = 0.01) association-test meta-analysis z-score map of the *social cognition* term (downloaded from Neurosynth; https://neurosynth.org/; Yarkoni et al., 2011) with the STAP (red outline) depicted. **B.** Same as **A.**, but for the *communication* term. **C.** Same as **A.**, but for the *autism* term.

Future studies examining the anterior STS, STAP, and pSTS relative to functional regions, as well as anatomical and functional connectivity, in individual hemispheres can provide further insight regarding the structural-functional heterogeneity of the STS. For example, Bodin et al. (2018) demonstrated that the STAP corresponds to voice-selective functional peaks in NT individuals, and whether this correspondence holds in ASD remains an open question. Future studies should test whether STS functional differences in ASD co-localize with the STAP, as well as extend to other portions of the STS.

### Novel anatomical and functional targets in ASD research

The present findings offer new anatomical and functional targets in ASD research, not just in the superior temporal cortex, but throughout the cerebral cortex more broadly. Here we focused on sulcal pits and laterality, and particularly the human-specific STAP. As sulcal pits are the first points of the cerebral cortex to pull in gestation as gyrencephalic brains begin to fold, sulcal pits provide developmental insight (Rakic, 1988; Zilles et al., 2013; Regis, 2005; Im et al., 2010; Borrell, 2018). Additionally, sulci emerge at different timepoints in which tertiary sulci emerging last in association cortices, and recent studies show that tertiary sulci are morphologically distinct in ASD individuals compared to NTs (Ammons et al., 2021; Brun et al., 2016; Nakamura et al., 2020; Ramos Benitez et al., 2024; <u>Willbrand et al., 2026a</u>). As all sulci have sulcal pits, and tertiary sulci have been largely overlooked due to the fact that they need to be manually defined (Miller, D’Esposito, and Weiner, 2021 for review), which is an arduous and time-consuming process, the present work lays a foundation for targeting sulcal pits in tertiary sulci as novel targets in future ASD research.

In addition to sulcal pits, recent research also highlights other fine-grained neuroanatomical features that are related to individual differences in development that could serve as new neuroanatomical targets in future ASD studies. For instance, a recent study showed that gyral crowns are related to face processing (<u>Willbrand et al., 2026b</u>). Specifically, gyral crowns in individuals with Developmental Prosopagnosia (DP) were morphologically different compared to NTs, were related to the size of functional regions, and predicted individual differences in cognition (<u>Willbrand et al., 2026b</u>). Another novel neuroanatomical developmental target is the cortical portion between a fractionated sulcus—referred to as *pli de passage* (Bodin et al., 2021; Mangin et al., 2019) or more modernly, gyral gaps (Borst et al., 2016; Bouhali et al., 2024; Cachia et al., 2018). Recently, changes in a gyral gap in the temporal cortex was related to reading ability (Borst et al., 2016; Bouhali et al., 2024; Cachia et al., 2018). Importantly, these changes were related to underlying changes in white matter properties, which highlights the relationship between, gyri, sulci, and network properties—specifically, short- and long-range connectivity. Long-range fibers have a bias to terminate in gyral crowns, while short-range fibers have a bias to terminate in the bottom, or fundi, of sulci (Cottaar et al., 2021; Reveley et al., 2015; Schilling et al., 2018). Thus, the present findings generate novel hypotheses for future ASD studies linking fine-grained (sulcal pits, gyral crowns, gyral gaps, etc.) novel neuroanatomical targets to functional regions, network organization, and individual differences in cognition. For example, both the aSTS and IFS showed a left-hemisphere bias in the present data. As previous research shows differences in the uncinate fasciculus, which connects inferior frontal and anterior temporal cortices, between ASD individuals and NTs (Andrews et al., 2021; Hung et al., 2023; Olivé et al., 2022; Yoon et al., 2026), future research can further explore this relationship linking sulcal morphology and underlying white matter between groups.

### Developmental trajectory of neuroanatomical structures as predictors for future studies

These findings also provide a foundation for novel hypotheses moving forward regarding developmental trajectories of cortical morphology. For example, as mentioned in the previous section, morphological features of sulci that emerge near term and continue to develop postnatally (tertiary sulci; Armstrong et al., 1995; Chi et al., 1977; Miller et al., 2021; Welker, 1990; Willbrand et al., 2022, 2023), have been found to differ in ASD (e.g., mid-fusiform sulcus length and thickness, Ammons et al., 2021; Ramos Benitez et al., 2024; paracingulate sulcus asymmetry, <u>Willbrand et al., 2026a</u>; decreased number of posterior orbital sulci, Nakamura et al., 2020). Nevertheless, previous and present findings show that early-forming neuroanatomical structures such as the STAP (Leroy et al., 2015) are not significantly different between individuals with ASD and NTs. This pattern may reflect the developmental trajectory of sulci, where sulcal features established earliest in utero appear morphologically similar in ASD, whereas later-developing sulci, which continue to develop after birth (Amiez et al., 2019; Cachia, A. et al., 2021; Dubois et al., 2019; Miller et al., 2021; Sanides, 1962, 1964; Voorhies et al., 2021; Willbrand et al., 2023; Welker, 1990), may show greater differences between groups. This interpretation is consistent with additional empirical studies in ASD. For instance, Brun et al. (2016) found that the sulcal pit depth of the ascending ramus of the lateral fissure in Broca’s region (Sprung-Much & Petrides, 2018) was reduced in ASD and correlated with individual differences in communication scores. Further, STS asymmetry (beyond the STAP) is detectable in fetuses as early as 26 weeks in gestation (Chi et al., 1977; Dubois et al., 2010; Habas et al., 2012). Finally, the STAP is preserved in the presence of situs inversus, Turner syndrome, and corpus callosum agenesis, suggesting that the development of STS asymmetry more broadly, and the STAP more focally, may be driven at least partly by genetic or molecular mechanisms operating somewhat independently of postnatal factors (Leroy et al., 2015). This raises the possibility that ASD-related neuroanatomical differences relative to NTs may be more apparent in the late prenatal and postnatal periods than in early gestation. Further studies of these later-developing sulci and sulcal pits in ASD could help clarify these similarities and differences. For example, longitudinal structural MRI studies of how sulci begin to diverge in ASD may help identify the factors that underlie sulcal formation, such as genetics, cellular mechanisms, or even biomechanical forces (Borrell, 2018; Dubois et al., 2008; Kroenke and Bayly, 2018).

In terms of limitations, participants were restricted to male individuals under 20 years old, limiting the generalizability of these findings to female and older adult populations. Leroy et al. (2015) reported that STAP asymmetry is greater in males than females in NT adults in a small sample (N=15), so whether the present findings extend to female participants in a large sample size remains untested.

## Conclusion

The present study empirically identifies the STAP as a stable neuroanatomical landmark in ASD in the largest sample size to date. Additionally, it provides clarity to previous findings that the STS is morphologically and functionally different between individuals with ASD and NTs. Specifically, these findings indicate that the differences between groups identified previously are likely driven by later emerging portions of the STS located anteriorly and posteriorly to the STAP, which emerges earlier in gestation. These findings underscore the value of studying individual sulcal morphology in ASD with a precision imaging approach, as well as support further exploration of the STAP in relation to longitudinal measurements of structure, function, and behavior across development, the lifespan, and in different clinical populations. We share average STAP labels for each group with the publication of this paper to guide such future research (**Fig. 1B** & **1C**; **Data Availability Statement**).

## Data Availability Statement

All data and original code used for the present project will be publicly available on GitHub upon publication (https://github.com/cnl-berkeley/stable_projects). Any additional information required to reanalyze the data reported in this paper is available from the corresponding author (Kevin Weiner) upon request.

## Ethics Approval Statement

Each contributing Autism Brain Imaging Data Exchange site confirmed that its local Institutional Review Board or ethics committee approved both the original data collection and the retrospective sharing of de-identified datasets. Contributing sites verified that all data were de-identified in accordance with HIPAA standards, including removal of all 18 protected health information identifiers and facial features from structural MRI scans.

## References

1. Allison, T., Puce, A., & McCarthy, G. (2000). Social perception from visual cues: Role of the STS region. Trends in Cognitive Sciences, 4(7), 267–278. 10.1016/S1364-6613(00)01501-1

2. Amiez, C., Sallet, J., Hopkins, W. D., Meguerditchian, A., Hadj-Bouziane, F., Ben Hamed, S., Wilson, C. R. E., Procyk, E., & Petrides, M. (2019). Sulcal organization in the medial frontal cortex provides insights into primate brain evolution. Nature Communications, 10, Article 3437. 10.1038/s41467-019-11347-x

3. Ammons, C. J., Winslett, M.-E., Bice, J., Patel, P., May, K. E., & Kana, R. K. (2021). The mid-fusiform sulcus in autism spectrum disorder: Establishing a novel anatomical landmark related to face processing. Autism Research, 14(1), 53–64. 10.1002/aur.2425

4. Andrews, D. S., Lee, J. K., Harvey, D. J., Waizbard-Bartov, E., Solomon, M., Rogers, S. J., Nordahl, C. W., & Amaral, D. G. (2021). A longitudinal study of white matter development in relation to changes in autism severity across early childhood. Biological Psychiatry, 89(5), 424–432. 10.1016/j.biopsych.2020.10.013

5. Armstrong, E., Schleicher, A., Omran, H., Curtis, M., & Zilles, K. (1995). The ontogeny of human gyrification. Cerebral Cortex, 5(1), 56–63. 10.1093/cercor/5.1.56

6. Auzias, G., Viellard, M., Takerkart, S., Villeneuve, N., Poinso, F., Da Fonséca, D., Girard, N., & Deruelle, C. (2014). Atypical sulcal anatomy in young children with autism spectrum disorder. NeuroImage: Clinical, 4, 593–603. 10.1016/j.nicl.2014.03.008

7. Beauchamp, M. S., Lee, K. E., Argall, B. D., & Martin, A. (2004). Integration of auditory and visual information about objects in superior temporal sulcus. Neuron, 41(5), 809–823. 10.1016/S0896-6273(04)00070-4

8. Belin, P., Zatorre, R., Lafaille, P. et al. Voice-selective areas in human auditory cortex. Nature 403, 309–312 (2000). 10.1038/35002078

9. Binder, J. R., Frost, J. A., Hammeke, T. A., Cox, R. W., Rao, S. M., & Prieto, T. (1997). Human brain language areas identified by functional magnetic resonance imaging. Journal of Neuroscience, 17(1), 353–362. 10.1523/JNEUROSCI.17-01-00353.1997

10. Boddaert, N., Chabane, N., Gervais, H., Good, C. D., Bourgeois, M., Plumet, M.-H., Barthélémy, C., Mouren, M.-C., Artiges, E., Samson, Y., Brunelle, F., Frackowiak, R. S. J., & Zilbovicius, M. (2004). Superior temporal sulcus anatomical abnormalities in childhood autism: A voxel-based morphometry MRI study. NeuroImage, 23(1), 364–369. 10.1016/j.neuroimage.2004.06.016

11. Bodin, C., Takerkart, S., Belin, P. et al. Anatomo-functional correspondence in the superior temporal sulcus. Brain Struct Funct 223, 221–232 (2018). 10.1007/s00429-017-1483-2

12. Bodin, C., Pron, A., Le Mao, M., Régis, J., Belin, P., & Coulon, O. (2021). Plis de passage in the superior temporal sulcus: Morphology and local connectivity. NeuroImage, 225, Article 117513. 10.1016/j.neuroimage.2020.117513

13. Borrell, V. (2018). How cells fold the cerebral cortex. Journal of Neuroscience, 38(4), 776–783. 10.1523/JNEUROSCI.1106-17.2017

14. Borst, G., Cachia, A., Tissier, C., Ahr, E., Simon, G., & Houdé, O. (2016). Early cerebral constraints on reading skills in school-age children: An MRI study. Mind, Brain, and Education, 10(1), 47–54. 10.1111/mbe.12098

15. Bouhali, F., Dubois, J., Hoeft, F., & Weiner, K. S. (2024). Unique longitudinal contributions of sulcal interruptions to reading acquisition in children. eLife. 10.7554/eLife.103007

16. Brass, M., Schmitt, R. M., Spengler, S., & Gergely, G. (2007). Investigating action understanding: Inferential processes versus action simulation. Current Biology, 17(24), 2117–2121. 10.1016/j.cub.2007.11.057

17. Brun, L., Auzias, G., Viellard, M., Villeneuve, N., Girard, N., Poinso, F., Da Fonseca, D., & Deruelle, C. (2016). Localized Misfolding Within Broca’s Area as a Distinctive Feature of Autistic Disorder. Biological psychiatry. Cognitive neuroscience and neuroimaging, 1(2), 160–168. 10.1016/j.bpsc.2015.11.003

18. Cachia, A., Roell, M., Mangin, J.-F., Sun, Z. Y., Jobert, A., Braga, L., Houdé, O., Dehaene, S., & Borst, G. (2018). How interindividual differences in brain anatomy shape reading accuracy. Brain Structure and Function, 223(2), 701–712. 10.1007/s00429-017-1516-x

19. Cachia, A., Borst, G., Jardri, R., Raznahan, A., Murray, G. K., Mangin, J.-F., & Plaze, M. (2021). Towards deciphering the fetal foundation of normal cognition and cognitive symptoms from sulcation of the cortex. Frontiers in Neuroanatomy, 15, Article 712862. 10.3389/fnana.2021.712862

20. Calvert, G. A., Hansen, P. C., Iversen, S. D., & Brammer, M. J. (2001). Detection of audio-visual integration sites in humans by application of electrophysiological criteria to the BOLD effect. NeuroImage, 14(2), 427–438. 10.1006/nimg.2001.0812

21. Cauda, F., Costa, T., Nani, A., Fava, L., Palermo, S., Bianco, F., Duca, S., Tatu, K., & Keller, R. (2017). Are schizophrenia, autistic, and obsessive spectrum disorders dissociable on the basis of neuroimaging morphological findings? A voxel-based meta-analysis. Autism Research, 10(6), 1079–1095. 10.1002/aur.1759

22. Chi JG, Dooling EC, Gilles FH. Gyral development of the human brain. Annals of Neurology. 1977:1:86–93.

23. Constantino, J. N., Davis, S. A., Todd, R. D., Schindler, M. K., Gross, M. M., Brophy, S. L., Metzger, L. M., Shoushtari, C. S., Splinter, R., & Reich, W. (2003). Validation of a brief quantitative measure of autistic traits: Comparison of the Social Responsiveness Scale with the Autism Diagnostic Interview–Revised. Journal of Autism and Developmental Disorders, 33(4), 427–433. 10.1023/A:1025014929212

24. Corbetta, M., & Shulman, G. L. (2002). Control of goal-directed and stimulus-driven attention in the brain. Nature Reviews Neuroscience, 3(3), 201–215. 10.1038/nrn755

25. Cottaar, M., Bastiani, M., Boddu, N., Glasser, M. F., Haber, S., van Essen, D. C., Sotiropoulos, S. N., & Jbabdi, S. (2021). Modelling white matter in gyral blades as a continuous vector field. NeuroImage, 227, Article 117693. 10.1016/j.neuroimage.2020.117693

26. Dale, A. M., Fischl, B., & Sereno, M. I. (1999). Cortical surface-based analysis: I. Segmentation and surface reconstruction. NeuroImage, 9(2), 179–194. 10.1006/nimg.1998.0395

27. Deen, B., Koldewyn, K., Kanwisher, N., & Saxe, R. (2015). Functional organization of social perception and cognition in the superior temporal sulcus. Cerebral Cortex, 25(11), 4596–4609. 10.1093/cercor/bhv111

28. de Vareilles, H., Rivière, D., Mangin, J. F., & Dubois, J. (2023). Development of cortical folds in the human brain: An attempt to review biological hypotheses, early neuroimaging investigations and functional correlates. Developmental Cognitive Neuroscience, 61, Article 101249. 10.1016/j.dcn.2023.101249

29. Di Martino, A., Yan, C.-G., Li, Q., Denio, E., Castellanos, F. X., Alaerts, K., Anderson, J. S., Assaf, M., Bookheimer, S. Y., Dapretto, M., Deen, B., Delmonte, S., Dinstein, I., Ertl-Wagner, B., Fair, D. A., Gallagher, L., Kennedy, D. P., Keown, C. L., Keysers, C., … Milham, M. P. (2014). The autism brain imaging data exchange: Towards a large-scale evaluation of the intrinsic brain architecture in autism. Molecular Psychiatry, 19(6), 659–667. 10.1038/mp.2013.78

30. Dubois, J., Benders, M., Cachia, A., Lazeyras, F., Ha-Vinh Leuchter, R., Sizonenko, S. V., Borradori-Tolsa, C., Mangin, J. F., & Hüppi, P. S. (2008). Mapping the early cortical folding process in the preterm newborn brain. Cerebral Cortex, 18(6), 1444–1454. 10.1093/cercor/bhm180

31. Dubois, J., Benders, M., Lazeyras, F., Borradori-Tolsa, C., Leuchter, R. H., Mangin, J. F., & Hüppi, P. S. (2010). Structural asymmetries of perisylvian regions in the preterm newborn. NeuroImage, 52(1), 32–42. 10.1016/j.neuroimage.2010.03.054

32. Dubois, J., Lefèvre, J., Angleys, H., Leroy, F., Fischer, C., Lebenberg, J., Dehaene-Lambertz, G., Borradori-Tolsa, C., Lazeyras, F., Hertz-Pannier, L., Mangin, J.-F., Hüppi, P. S., & Germanaud, D. (2019). The dynamics of cortical folding waves and prematurity-related deviations revealed by spatial and spectral analysis of gyrification. NeuroImage, 185, 934–946. 10.1016/j.neuroimage.2018.03.005

33. Ecker, C., Ginestet, C., Feng, Y., Johnston, P., Lombardo, M. V., Lai, M.-C., Suckling, J., Palaniyappan, L., Daly, E., Murphy, C. M., Williams, S. C., Bullmore, E. T., Baron-Cohen, S., Brammer, M., Murphy, D. G. M., & MRC AIMS Consortium. (2013). Brain surface anatomy in adults with autism: The relationship between surface area, cortical thickness, and autistic symptoms. JAMA Psychiatry, 70(1), 59–70. 10.1001/jamapsychiatry.2013.265

34. Fan, X.-R., He, Y., Wang, Y.-S., Li, L., Duan, X., & Zuo, X.-N. (2025). Profiling brain morphology for autism spectrum disorder with two cross-culture large-scale consortia. Communications Biology, 8, Article 1157. 10.1038/s42003-025-08573-z

35. Fedorenko, E., Nieto-Castañón, A., & Kanwisher, N. (2012). Lexical and syntactic representations in the brain: An fMRI investigation with multi-voxel pattern analyses. Neuropsychologia, 50(4), 499–513. 10.1016/j.neuropsychologia.2011.09.014

36. Feng, Y., Zhao, W., Li, Y., Yin, Q., Wang, X., Huang, X., Li, L., Shan, X., Hu, W., Ming, Y., Wang, P., Xiao, J., Chen, H., & Duan, X. (2025). Diffusion trajectory of atypical morphological development in autism spectrum disorder. Communications Biology, 8, Article 1476. 10.1038/s42003-025-08874-3

37. Fischl, B., Sereno, M. I., & Dale, A. M. (1999). Cortical surface-based analysis: II. Inflation, flattening, and a surface-based coordinate system. NeuroImage, 9(2), 195–207. 10.1006/nimg.1998.0396

38. Fletcher, P. C., Happé, F., Frith, U., Baker, S. C., Dolan, R. J., Frackowiak, R. S. J., & Frith, C. D. (1995). Other minds in the brain: A functional imaging study of “theory of mind” in story comprehension. Cognition, 57(2), 109–128. 10.1016/0010-0277(95)00692-R

39. Fornito, A., Yücel, M., Wood, S. J., Proffitt, T., McGorry, P. D., Velakoulis, D., & Pantelis, C. (2006). Morphology of the paracingulate sulcus and executive cognition in schizophrenia. Schizophrenia Research, 88(1–3), 192–197. 10.1016/j.schres.2006.06.034

40. Gallagher, H. L., Happé, F., Brunswick, N., Fletcher, P. C., Frith, U., & Frith, C. D. (2000). Reading the mind in cartoons and stories: An fMRI study of “theory of mind” in verbal and nonverbal tasks. Neuropsychologia, 38(1), 11–21. 10.1016/S0028-3932(99)00053-6

41. Garrison, J. R., Fernyhough, C., McCarthy-Jones, S., Haggard, M., The Australian Schizophrenia Research Bank, & Simons, J. S. (2015). Paracingulate sulcus morphology is associated with hallucinations in the human brain. Nature Communications, 6, Article 8956. 10.1038/ncomms9956

42. Gervais, H., Belin, P., Boddaert, N., Leboyer, M., Coez, A., Sfaello, I., Barthélémy, C., Brunelle, F., Samson, Y., & Zilbovicius, M. (2004). Abnormal cortical voice processing in autism. Nature Neuroscience, 7(8), 801–802. 10.1038/nn1291

43. Gotham, K., Pickles, A., & Lord, C. (2009). Standardizing ADOS scores for a measure of severity in autism spectrum disorders. Journal of autism and developmental disorders, 39(5), 693–705. 10.1007/s10803-008-0674-3

44. Grossman, E., Donnelly, M., Price, R., Pickens, D., Morgan, V., Neighbor, G., & Blake, R. (2000). Brain areas involved in perception of biological motion. Journal of Cognitive Neuroscience, 12(5), 711–720. 10.1162/089892900562417

45. Guo, Z., Tang, X., Xiao, S., Yan, H., Sun, S., Yang, Z., Huang, L., Chen, Z., & Wang, Y. (2024). Systematic review and meta-analysis: Multimodal functional and anatomical neural alterations in autism spectrum disorder. Molecular Autism, 15, Article 16. 10.1186/s13229-024-00593-6

46. Habas, P. A., Scott, J. A., Roosta, A., Rajagopalan, V., Kim, K., Rousseau, F., Barkovich, A. J., Glenn, O. A., & Studholme, C. (2012). Early folding patterns and asymmetries of the normal human brain detected from in utero MRI. Cerebral Cortex, 22(1), 13–25. 10.1093/cercor/bhr053

47. Hastings, W. L., III, Willbrand, E. H., Kelly, J. P., Washington, S. T., Tameilau, P., Sathishkumar, R. N., Maboudian, S. A., Parker, B. J., Elliott, M. V., Johnson, S. L., & Weiner, K. S. (2024). Emotion-related impulsivity is related to orbitofrontal cortical sulcation. Cortex, 181, 140–154. 10.1016/j.cortex.2024.08.009

48. Haxby, J. V., Hoffman, E. A., & Gobbini, M. I. (2000). The distributed human neural system for face perception. Trends in Cognitive Sciences, 4(6), 223–233. 10.1016/s1364-6613(00)01482-0

49. Hung, Y., Dallenbach, N. T., Green, A., Gaillard, S., Capella, J., Hoskova, B., Vater, C. H., Cooper, E., Rudberg, N., Takahashi, A., Gabrieli, J. D. E., & Joshi, G. (2023). Distinct and shared white matter abnormalities when ADHD is comorbid with ASD: A preliminary diffusion tensor imaging study. Psychiatry Research, 320, Article 115039. 10.1016/j.psychres.2022.115039

50. Im, K., Jo, H. J., Mangin, J.-F., Evans, A. C., Kim, S. I., & Lee, J.-M. (2010). Spatial distribution of deep sulcal landmarks and hemispherical asymmetry on the cortical surface. Cerebral Cortex, 20(3), 602–611. 10.1093/cercor/bhp127

51. Keown, C. L., Shih, P., Nair, A., Peterson, N., Mulvey, M. E., & Müller, R.-A. (2013). Local functional overconnectivity in posterior brain regions is associated with symptom severity in autism spectrum disorders. Cell Reports, 5(3), 567–572. 10.1016/j.celrep.2013.10.003

52. Kroenke, C. D., & Bayly, P. V. (2018). How forces fold the cerebral cortex. Journal of Neuroscience, 38(4), 767–775. 10.1523/JNEUROSCI.1105-17.2017

53. Le Guen, Y., Leroy, F., Auzias, G., Rivière, D., Grigis, A., Mangin, J. F., Coulon, O., Dehaene-Lambertz, G., & Frouin, V. (2018). The chaotic morphology of the left superior temporal sulcus is genetically constrained. NeuroImage, 174, 297–307. 10.1016/j.neuroimage.2018.03.046

54. Le Guen, Y., Leroy, F., Philippe, C., IMAGEN Consortium, Mangin, J. F., Dehaene-Lambertz, G., & Frouin, V. (2020). Enhancer locus in ch14q23.1 modulates brain asymmetric temporal regions involved in language processing. Cerebral Cortex, 30(10), 5322–5332. 10.1093/cercor/bhaa112

55. Leroy, F., Cai, Q., Bogart, S. L., Dubois, J., Coulon, O., Monzalvo, K., Fischer, C., Glasel, H., Van der Haegen, L., Bénézit, A., Lin, C.-P., Kennedy, D. N., Ihara, A. S., Hertz-Pannier, L., Moutard, M.-L., Poupon, C., Brysbaert, M., Roberts, N., Hopkins, W. D., … Dehaene-Lambertz, G. (2015). New human-specific brain landmark: The depth asymmetry of superior temporal sulcus. Proceedings of the National Academy of Sciences, 112(4), 1208–1213. 10.1073/pnas.1412389112

56. Levitt, J. G., Blanton, R. E., Smalley, S., Thompson, P. M., Guthrie, D., McCracken, J. T., Sadoun, T., Heinichen, L., & Toga, A. W. (2003). Cortical sulcal maps in autism. Cerebral Cortex, 13(7), 728–735. 10.1093/cercor/13.7.728

57. Li, C., Park, H., Awasthi, J., Rolison, M., Li, M., Scheinost, D., Chawarska, K., & Hampson, M. (2025). Disrupted resting-state functional connectivity in the social visual pathway in children with autism spectrum disorder. Autism Research, 18(5), 1024–1036. 10.1002/aur.70037

58. Lohmann, G., von Cramon, D. Y., & Colchester, A. C. F. (2008). Deep sulcal landmarks provide an organizing framework for human cortical folding. Cerebral Cortex, 18(6), 1415–1420. 10.1093/cercor/bhm174

59. Lord, C., Rutter, M., DiLavore, P. C., Risi, S., Gotham, K., & Luyster, R. J. (2012). Autism Diagnostic Observation Schedule, Second Edition (ADOS-2) Manual. Los Angeles, CA: Western Psychological Services

60. Maboudian, S. A., Willbrand, E. H., Kelly, J. P., Jagust, W. J., & Weiner, K. S., for the Alzheimer’s Disease Neuroimaging Initiative. (2024). Defining overlooked structures reveals new associations between the cortex and cognition in aging and Alzheimer’s disease. Journal of Neuroscience, 44(16), Article e1714232024. 10.1523/JNEUROSCI.1714-23.2024

61. Mangin, J.-F., Le Guen, Y., Labra, N., Grigis, A., Frouin, V., Guevara, M., Fischer, C., Rivière, D., Hopkins, W. D., Régis, J., & Sun, Z. Y. (2019). “Plis de passage” deserve a role in models of the cortical folding process. Brain Topography, 32(6), 1035–1048. 10.1007/s10548-019-00734-8

62. Meng, Y., Li, G., Lin, W., Gilmore, J. H., & Shen, D. (2014). Spatial distribution and longitudinal development of deep cortical sulcal landmarks in infants. NeuroImage, 100, 206–218. 10.1016/j.neuroimage.2014.06.004

63. Miller, J. A., D’Esposito, M., & Weiner, K. S. (2021). Using tertiary sulci to map the “cognitive globe” of prefrontal cortex. Journal of Cognitive Neuroscience, 33(9), 1698–1715. 10.1162/jocn_a_01696

64. Minshew, N. J., & Keller, T. A. (2010). The nature of brain dysfunction in autism: functional brain imaging studies. Current opinion in neurology, 23(2), 124–130. 10.1097/WCO.0b013e32833782d4

65. Nakamura, M., Nestor, P. G., & Shenton, M. E. (2020). Orbitofrontal Sulcogyral Pattern as a Transdiagnostic Trait Marker of Early Neurodevelopment in the Social Brain. Clinical EEG and Neuroscience. 10.1177/1550059420904180

66. Natu, V. S., Arcaro, M. J., Barnett, M. A., Gomez, J., Livingstone, M., Grill-Spector, K., & Weiner, K. S. (2021). Sulcal depth in the medial ventral temporal cortex predicts the location of a place-selective region in macaques, children, and adults. Cerebral Cortex, 31(1), 48–61. 10.1093/cercor/bhaa203

67. Nordahl, C. W., Dierker, D., Mostafavi, I., Schumann, C. M., Rivera, S. M., Amaral, D. G., & Van Essen, D. C. (2007). Cortical folding abnormalities in autism revealed by surface-based morphometry. Journal of Neuroscience, 27(43), 11725–11735. 10.1523/JNEUROSCI.0777-07.2007

68. Ohnishi, T., Matsuda, H., Hashimoto, T., Kunihiro, T., Nishikawa, M., Uema, T., & Sasaki, M. (2000). Abnormal regional cerebral blood flow in childhood autism. Brain, 123(9), 1838–1844. 10.1093/brain/123.9.1838

69. Olivé, G., Slušná, D., Vaquero, L., Muchart-López, J., Rodríguez-Fornells, A., & Hinzen, W. (2022). Structural connectivity in ventral language pathways characterizes non-verbal autism. Brain Structure and Function, 227(5), 1817–1829. 10.1007/s00429-022-02474-1

70. Ong, L. T., & Fan, S. W. D. (2023). Morphological and functional changes of cerebral cortex in autism spectrum disorder. Innovations in Clinical Neuroscience, 20(10–12), 40–47.

71. Parker, B.J., Voorhies, W.I., Jiahui, G. et al. Hominoid-specific sulcal variability is related to face perception ability. Brain Struct Funct 228, 677–685 (2023). 10.1007/s00429-023-02611-4

72. Pelphrey, K. A., Morris, J. P., Michelich, C. R., Allison, T., & McCarthy, G. (2005). Functional anatomy of biological motion perception in posterior temporal cortex: An fMRI study of eye, mouth and hand movements. Cerebral Cortex, 15(12), 1866–1876. 10.1093/cercor/bhi064

73. Pitcher, D., Dilks, D. D., Saxe, R. R., Triantafyllou, C., & Kanwisher, N. (2011). Differential selectivity for dynamic versus static information in face-selective cortical regions. NeuroImage, 56(4), 2356–2363. 10.1016/j.neuroimage.2011.03.067

74. Pitcher, D., & Ungerleider, L. G. (2021). Evidence for a third visual pathway specialized for social perception. Trends in Cognitive Sciences, 25(2), 100–110. 10.1016/j.tics.2020.11.006

75. Puce, A. (2024). From motion to emotion: Visual pathways and potential interconnections. Journal of Cognitive Neuroscience, 36(12), 2594–2617. 10.1162/jocn_a_02141

76. Rakic, P. (1988). Specification of cerebral cortical areas. Science, 241(4862), 170–176. 10.1126/science.3291116

77. Ramos Benitez, J., Kannan, S., Hastings, W. L., Parker, B. J., Willbrand, E. H., & Weiner, K. S. (2024). Ventral temporal and posteromedial sulcal morphology in autism spectrum disorder. Neuropsychologia, 195, Article 108786. 10.1016/j.neuropsychologia.2024.108786

78. Régis, J., Mangin, J.-F., Ochiai, T., Frouin, V., Rivière, D., Cachia, A., Tamura, M., & Samson, Y. (2005). “Sulcal root” generic model: a hypothesis to overcome the variability of the human cortex folding patterns. Neurologia medico-chirurgica, 45(1), 1–17.

79. Reveley, C., Seth, A. K., Pierpaoli, C., Silva, A. C., Yu, D., Saunders, R. C., Leopold, D. A., & Ye, F. Q. (2015). Superficial white matter fiber systems impede detection of long-range cortical connections in diffusion MR tractography. Proceedings of the National Academy of Sciences of the United States of America, 112(21), E2820–E2828. 10.1073/pnas.1418198112

80. Sanides, F. (1962). *Die Architektonik des menschlichen Stirnhirns und angrenzender Felder* (Monographien aus dem Gesamtgebiete der Neurologie und Psychiatrie, Heft 98). Springer.

81. Sanides, F. (1964). Structure and function of the human frontal lobe. Neuropsychologia, 2(3), 209–219. 10.1016/0028-3932(64)90005-3

82. Saxe, R., & Kanwisher, N. (2003). People thinking about thinking people: The role of the temporo-parietal junction in “theory of mind.“ NeuroImage, 19(4), 1835–1842. 10.1016/S1053-8119(03)00230-1

83. Schilling, K. G., Gao, Y., Janve, V., Stepniewska, I., Landman, B. A., & Anderson, A. W. (2018). Confirmation of a gyral bias in diffusion MRI fiber tractography. Human Brain Mapping, 39(3), 1449–1466. 10.1002/hbm.23936

84. Shih, P., Keehn, B., Oram, J. K., Leyden, K. M., Keown, C. L., & Müller, R.-A. (2011). Functional differentiation of posterior superior temporal sulcus in autism: A functional connectivity magnetic resonance imaging study. Biological Psychiatry, 70(3), 270–277. 10.1016/j.biopsych.2011.03.040

85. Sprung-Much, T., & Petrides, M. (2018). Morphological patterns and spatial probability maps of two defining sulci of the posterior ventrolateral frontal cortex of the human brain: The sulcus diagonalis and the anterior ascending ramus of the lateral fissure. Brain Structure and Function, 223(9), 4125–4152. 10.1007/s00429-018-1733-y

86. Stickel, S., Weismann, P., Kellermann, T., Regenbogen, C., Habel, U., Freiherr, J., & Chechko, N. (2019). Audio–visual and olfactory–visual integration in healthy participants and subjects with autism spectrum disorder. Human Brain Mapping, 40(15), 4470–4486. 10.1002/hbm.24715

87. Taylor, K. I., Moss, H. E., Stamatakis, E. A., & Tyler, L. K. (2006). Binding crossmodal object features in perirhinal cortex. Proceedings of the National Academy of Sciences, 103(21), 8239–8244. 10.1073/pnas.0509704103

88. Tezcan, M. E., Ataş, A. E., Göncüoğlu, A., Ekici, F., & Kozanhan, B. (2025). Are olfactory bulb volume decreases and olfactory sulcus deepening associated with atypical sensory behaviours in children with autism spectrum disorders? Journal of Psychiatric Research, 184, 176–186. 10.1016/j.jpsychires.2025.02.063

89. Tung, S. S., Yan, X., Fascendini, B., Tyagi, C., Martinez Reyes, C., Ducre, K., Perez, K., Allen, A., Horenziak, J., Wu, H., Keil, B., Natu, V. S., & Grill-Spector, K. (2026). How do infant brains fold? Sulcal deepening is linked to development of sulcal span, thickness, curvature, and microstructure. Communications Biology, 9, Article 258. 10.1038/s42003-026-09536-8

90. Van Essen, D. C. (1997). A tension-based theory of morphogenesis and compact wiring in the central nervous system. Nature, 385(6614), 313–318. 10.1038/385313a0

91. Vander Wyk, B. C., Hudac, C. M., Carter, E. J., Sobel, D. M., & Pelphrey, K. A. (2009). Action understanding in the superior temporal sulcus region. Psychological Science, 20(6), 771–777. 10.1111/j.1467-9280.2009.02359.x

92. Vigneau, M., Beaucousin, V., Hervé, P.-Y., Duffau, H., Crivello, F., Houdé, O., Mazoyer, B., & Tzourio-Mazoyer, N. (2006). Meta-analyzing left hemisphere language areas: Phonology, semantics, and sentence processing. NeuroImage, 30(4), 1414–1432. 10.1016/j.neuroimage.2005.11.002

93. Voorhies, W. I., Miller, J. A., Yao, J. K., Bunge, S. A., & Weiner, K. S. (2021). Cognitive insights from tertiary sulci in prefrontal cortex. Nature Communications, 12, 5122. 10.1038/s41467-021-25162-w

94. Watanabe, H., Nakamura, M., Ohno, T., Itahashi, T., Tanaka, E., Ohta, H., Yamada, T., Kanai, C., Iwanami, A., & Kato, N. (2014). Altered orbitofrontal sulcogyral patterns in adult males with high-functioning autism spectrum disorders. Social Cognitive and Affective Neuroscience, 9(4), 520–528. 10.1093/scan/nst016

95. Welker W. Why does cerebral cortex fissure and fold? In: Jones EG, Peters A, editors. Cerebral cortex: comparative structure and evolution of cerebral cortex, part II. Boston, MA: Springer US; 1990. pp. 3–136.

96. Willbrand, E. H., Parker, B. J., Voorhies, W. I., Miller, J. A., Lyu, I., Hallock, T., Aponik-Gremillion, L., Koslov, S. R., Alzheimer’s Disease Neuroimaging Initiative, Bunge, S. A., Foster, B. L., & Weiner, K. S. (2022). Uncovering a tripartite landmark in posterior cingulate cortex. Science Advances, 8(36), Article eabn9516. 10.1126/sciadv.abn9516

97. Willbrand, E. H., Ferrer, E., Bunge, S. A., & Weiner, K. S. (2023). Development of human lateral prefrontal sulcal morphology and its relation to reasoning performance. Journal of Neuroscience, 43(14), 2552–2567. 10.1523/JNEUROSCI.1745-22.2023

98. Willbrand, E. H., Jackson, S., Chen, S., Hathaway, C. B., Voorhies, W. I., Bunge, S. A., & Weiner, K. S. (2024). Sulcal variability in anterior lateral prefrontal cortex contributes to variability in reasoning performance among young adults. Brain Structure and Function, 229(2), 387–402. 10.1007/s00429-023-02734-8

99. Willbrand, E. H., Martinez, E., Ludwig, J. J., Maboudian, S. A., & Weiner, K. S. (2026). Anterior cingulate folding pattern is altered in autism spectrum disorder. Cerebral Cortex, 36(3), bhag012. 10.1093/cercor/bhag012

100. Willbrand, E. H., Kelly, J. P., Chen, X., Zhen, Z., Jiahui, G., Duchaine, B., & Weiner, K. S. (2026). Gyral crowns contribute to the cortical infrastructure of human face processing. Journal of Neuroscience, 46(2), Article e0562252025. 10.1523/JNEUROSCI.0562-25.2025

101. Willbrand, E. H., Tsai, Y.-H., Gagnant, T., & Weiner, K. S. (2026). Updating the sulcal landscape of the human lateral parieto-occipital junction provides anatomical, functional, and cognitive insights. eLife, 12, Article RP90451. 10.7554/eLife.90451

102. Willbrand, E. H., Maboudian, S. A., Elliott, M. V., Kellerman, G. M., Johnson, S. L., & Weiner, K. S. (2026). Variable presence of an evolutionarily new brain structure is related to trait impulsivity. Biological Psychiatry: Cognitive Neuroscience and Neuroimaging, 11(1), 8–16. 10.1016/j.bpsc.2024.11.015

103. Wu, S.-C. J., de Vareilles, H., Mitchell, S. C., Al-Manea, A., Garrison, J., Mamalakis, M., Simons, J. S., Cachia, A., Mangin, J.-F., Nerland, S., Mørch-Johnsen, L., Agartz, I., Suckling, J., & Murray, G. K. (2026). Reassessing asymmetry reduction in psychosis: Cingulate folding and gyrification covariance in patients with auditory hallucinations. Schizophrenia Bulletin, 52(2), sbaf086. 10.1093/schbul/sbaf086

104. Yarkoni, T., Poldrack, R. A., Nichols, T. E., Van Essen, D. C., & Wager, T. D. (2011). Large-scale automated synthesis of human functional neuroimaging data. Nature Methods, 8(8), 665–670. 10.1038/nmeth.1635

105. Yoon, C. D., Alexander, A. L., Travers, B. G., Lainhart, J. E., & Dean, D. C., III. (2026). Widespread alterations in white matter microstructure in autism: A multilevel meta-analysis. Biological Psychiatry: Cognitive Neuroscience and Neuroimaging. Advance online publication. 10.1016/j.bpsc.2026.08.002

106. Yücel, M., Stuart, G. W., Maruff, P., Velakoulis, D., Crowe, S. F., Savage, G., & Pantelis, C. (2002). Hemispheric and gender-related differences in the gross morphology of the anterior cingulate/paracingulate cortex in normal volunteers: An MRI morphometric study. Biological Psychiatry, 52(1), 15–23. 10.1016/S0006-3223(02)01312-4

107. Zeidan, J., Fombonne, E., Scorah, J., Ibrahim, A., Durkin, M. S., Saxena, S., Yusuf, A., Shih, A., & Elsabbagh, M. (2022). Global prevalence of autism: A systematic review update. Autism research : official journal of the International Society for Autism Research, 15(5), 778–790. 10.1002/aur.2696

108. Zilbovicius, M., Boddaert, N., Belin, P., Poline, J.-B., Remy, P., Mangin, J.-F., Thivard, L., Barthélémy, C., & Samson, Y. (2000). Temporal lobe dysfunction in childhood autism: A PET study. American Journal of Psychiatry, 157(12), 1988–1993. 10.1176/appi.ajp.157.12.1988

109. Zilbovicius, M., Meresse, I., Chabane, N., Brunelle, F., Samson, Y., & Boddaert, N. (2006). Autism, the superior temporal sulcus and social perception. Trends in Neurosciences, 29(7), 359–366. 10.1016/j.tins.2006.06.004

110. Zilles, K., Palomero-Gallagher, N., & Amunts, K. (2013). Development of cortical folding during evolution and ontogeny. Trends in Neurosciences, 36(5), 275–284.

