## Supplementary Materials for "The superior temporal asymmetrical pit remains asymmetric in autism spectrum disorder"

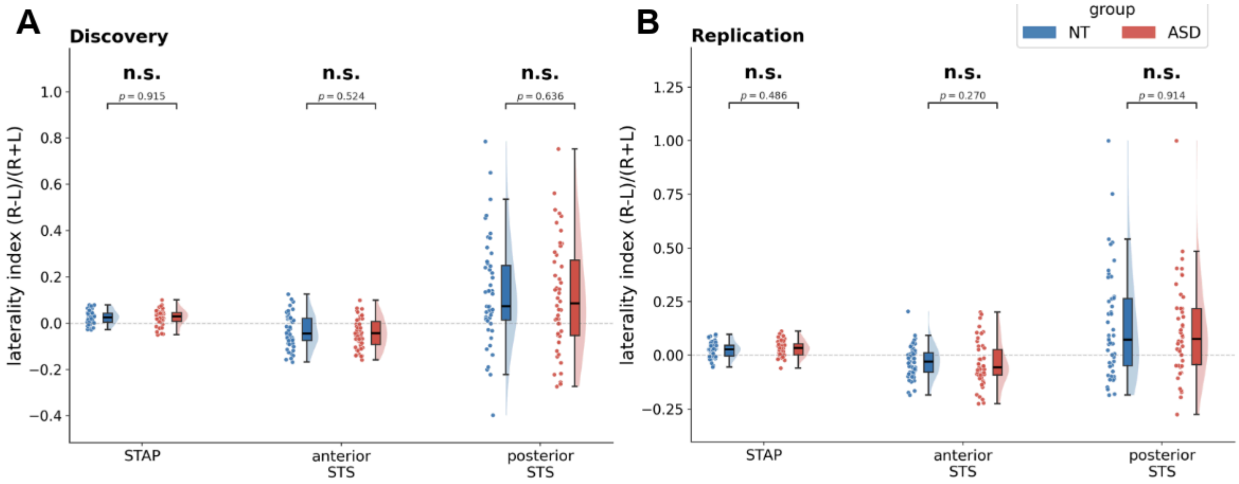


**Figure S1. STAP and STS laterality is similar between ASD and NT individuals. A.** Mean STAP (left), anterior STS (middle), and posterior STS (right) laterality (calculated via laterality index; LI) compared between NT (blue) and ASD (red) individuals in the discovery dataset. Individual dots represent individual subjects; dashed line at 0 STAP LI indicates symmetric hemispheres. **B.** Same as **A.**, analyzing the independent replication dataset. n.s. $p > .05$

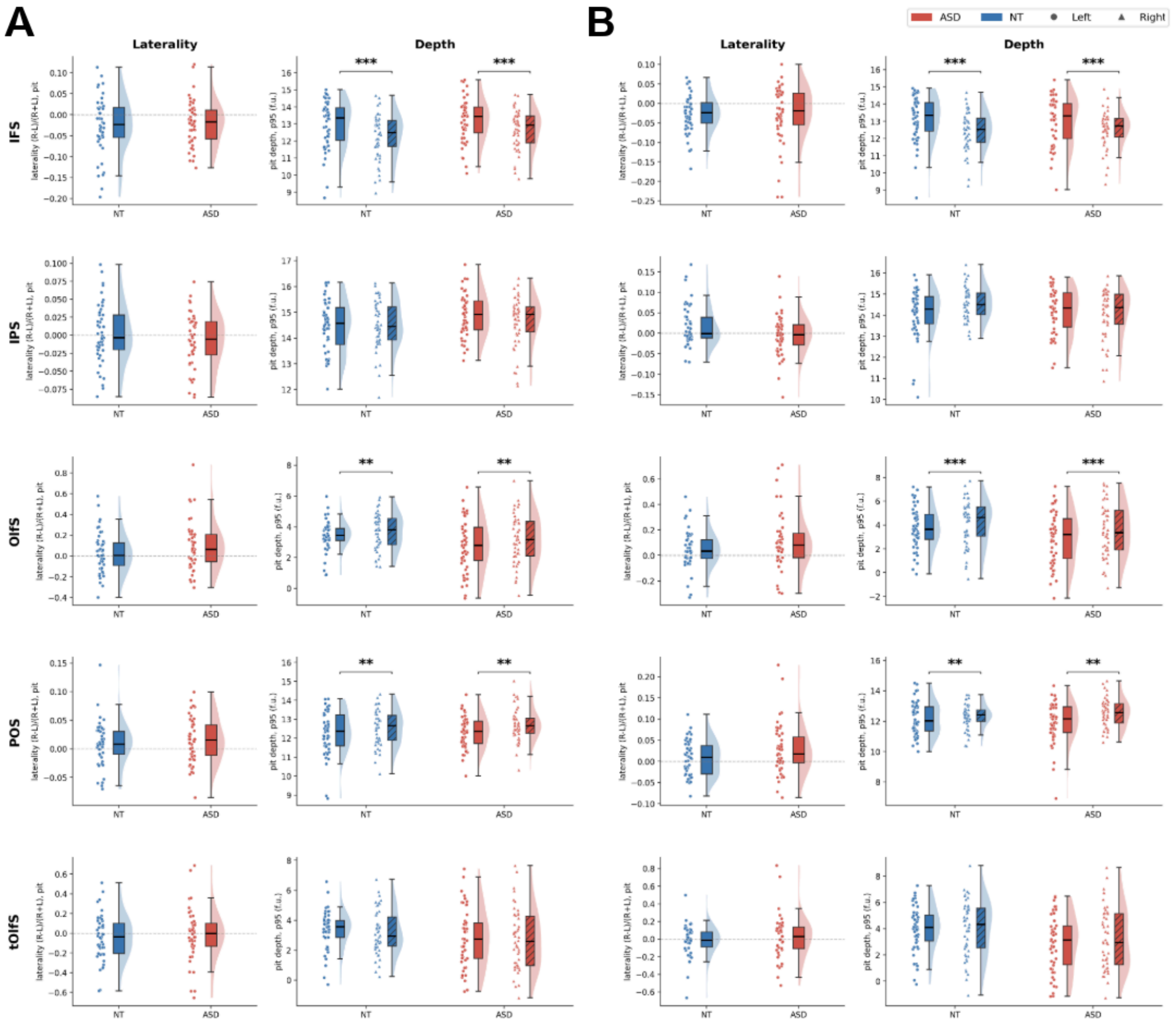


**Figure S2. No group differences between NT and ASD individuals for additional sulci tested outside the STS. A.** Raincloud plots visualizing sulci in NT (blue) and ASD (red) individuals within the discovery dataset. *Left*: LI; *Right*: left and right hemisphere pit mean depth. Sulci are listed in the following order from top to bottom rows: IFS, IPS, OlfS, POS, tOlfS. **B.** Same as **A.**, but for the independent replication dataset. f.u. = FreeSurfer units (Dale et al., 1999; Fischl et al., 1999), ***p < .001, **p < .01, n.s. p > .05. In the discovery sample (**Fig. S2A**), there were no group differences in pit depth (group main effect: all p > .06; group × hemisphere: all p > .23) or pit LI (all p > .18) for any of the additional sulci. The replication sample (**Fig. S2B**) similarly showed no group differences in pit LI (all p > .09) or pit depth (all other p > .05), with the exception of a main group effect for the tOlfS, with deeper pits in NT than ASD participants ($p = .016$), and a group × hemisphere interaction for the IPS ($p = .030$). However, these results were not replicated in the discovery sample (tOlfs main group effect: $p = .12$; IPS group × hemisphere: $p = .24$).

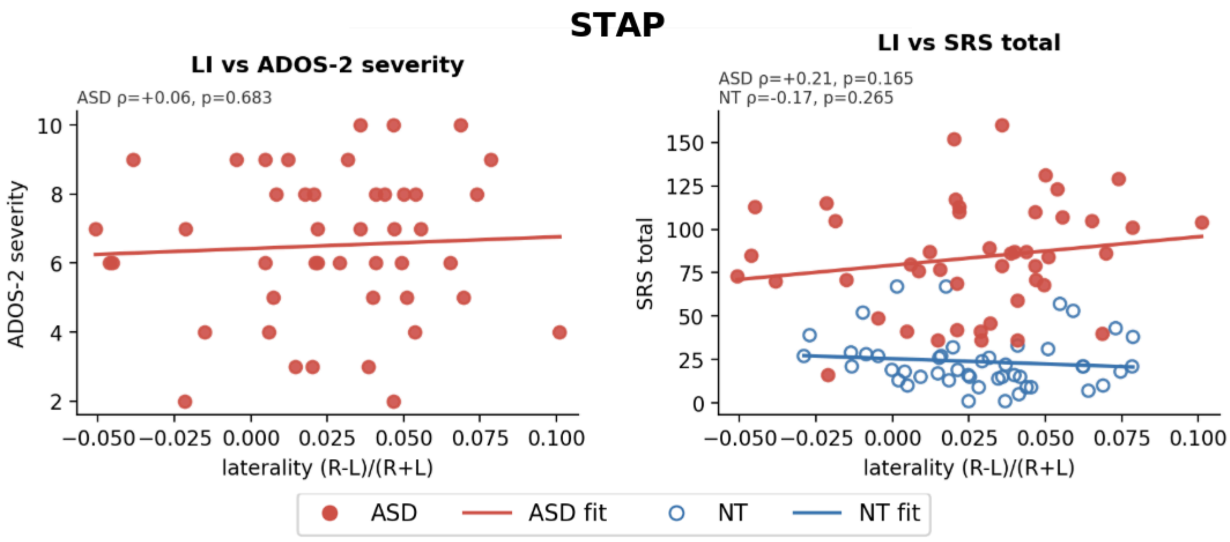


**Figure S3. STAP Laterality does not correlate with ADOS-2 or SRS scores.** Scatterplots visualizing the effects of STAP laterality (measured by the laterality index; LI) on ADOS-2 (left) and SRS (right) severity scores. Individual dots represent individual participants (ASD: red, filled; NT: blue, hollow). Solid red (ASD) and blue (NT) lines represent lines of best fit. ρ = Spearman's correlation coefficient. Not significant $p > .05$

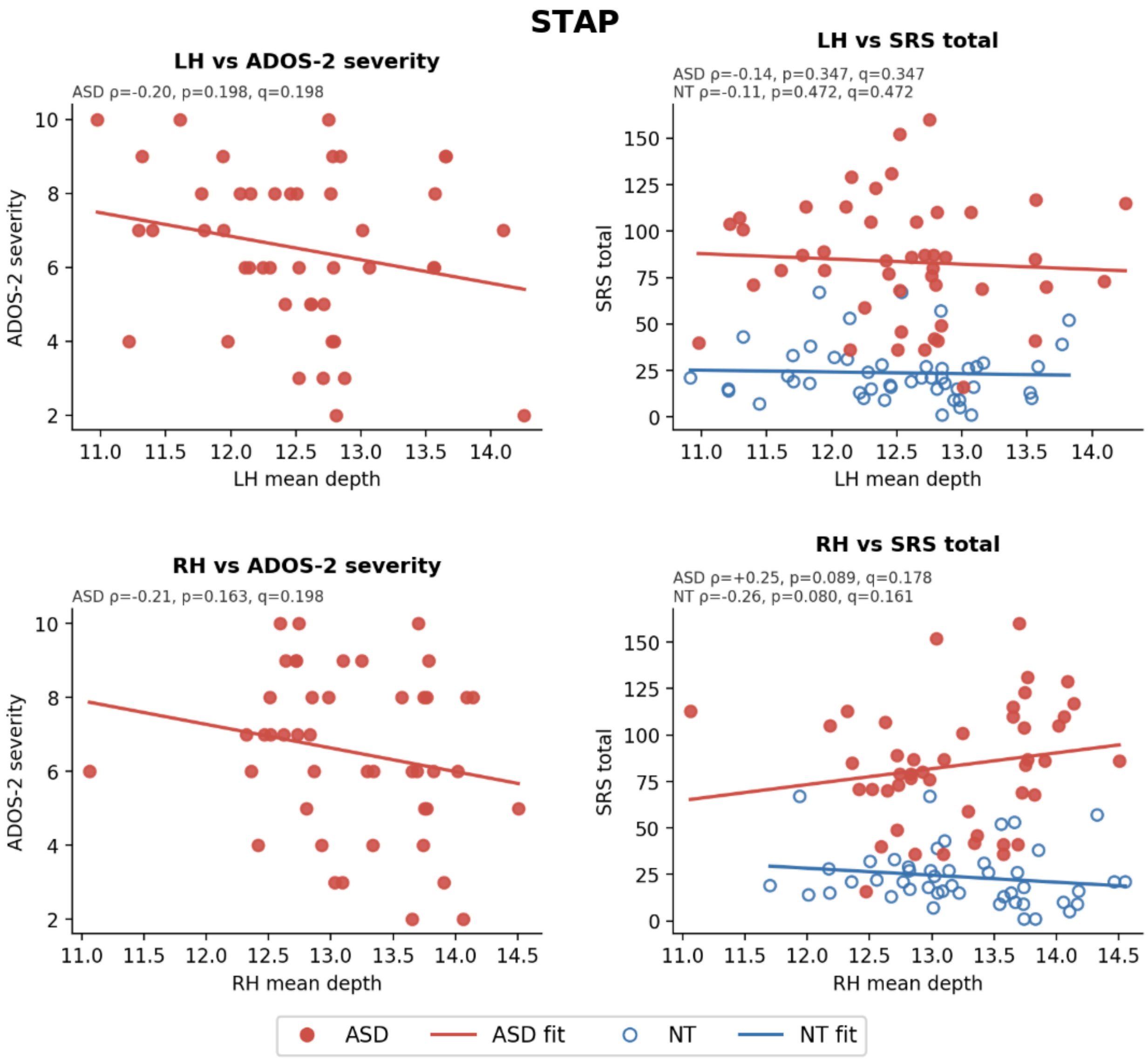


**Figure S4. Mean depth of the STAP does not correlate with ASD behavioral metrics.** Scatterplots visualizing the effects of STAP mean depth (measured in FreeSurfer Units; Dale et al., 1999; Fischl et al., 1999) on ADOS-2 (left column) and SRS (right column) severity scores. Individual dots represent individual participants (ASD: red, filled; NT: blue, hollow). Solid red (ASD) and blue (NT) lines represent lines of best fit.

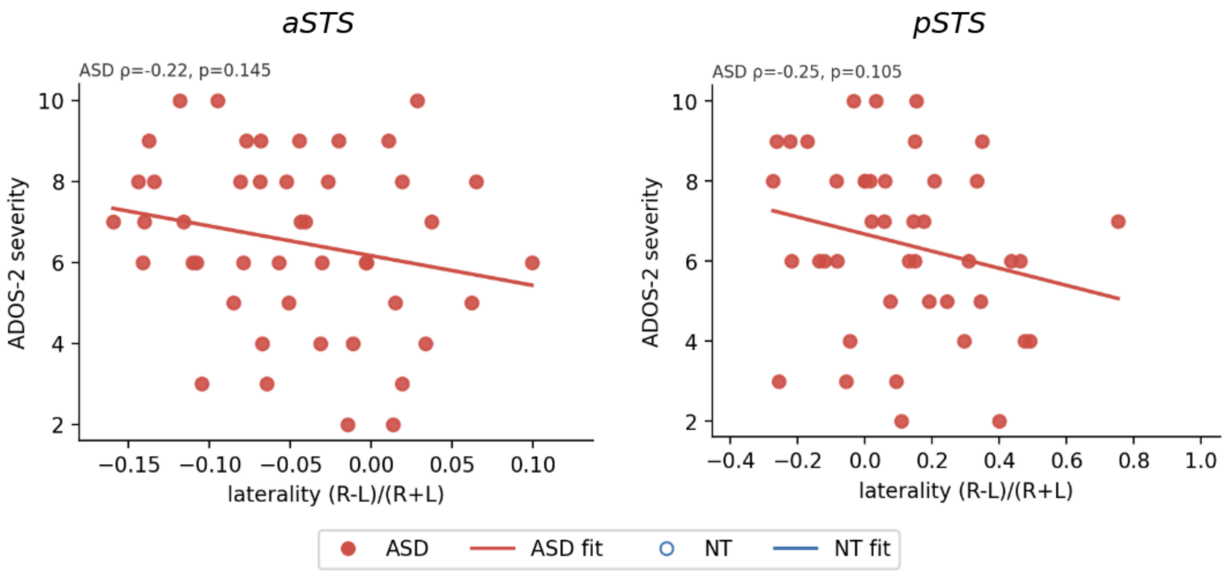


**Figure S5. Laterality of the anterior and posterior STS does not correlate with ADOS-2 severity scores.** Scatterplots visualizing the effects of anterior and posterior STS laterality (measured by the laterality index; LI) on ADOS-2 severity scores. Individual dots represent individual participants (ASD: red, filled; NT: blue, hollow). Solid red (ASD) lines represent lines of best fit. ρ = Spearman's correlation coefficient. Not significant $p > .05$

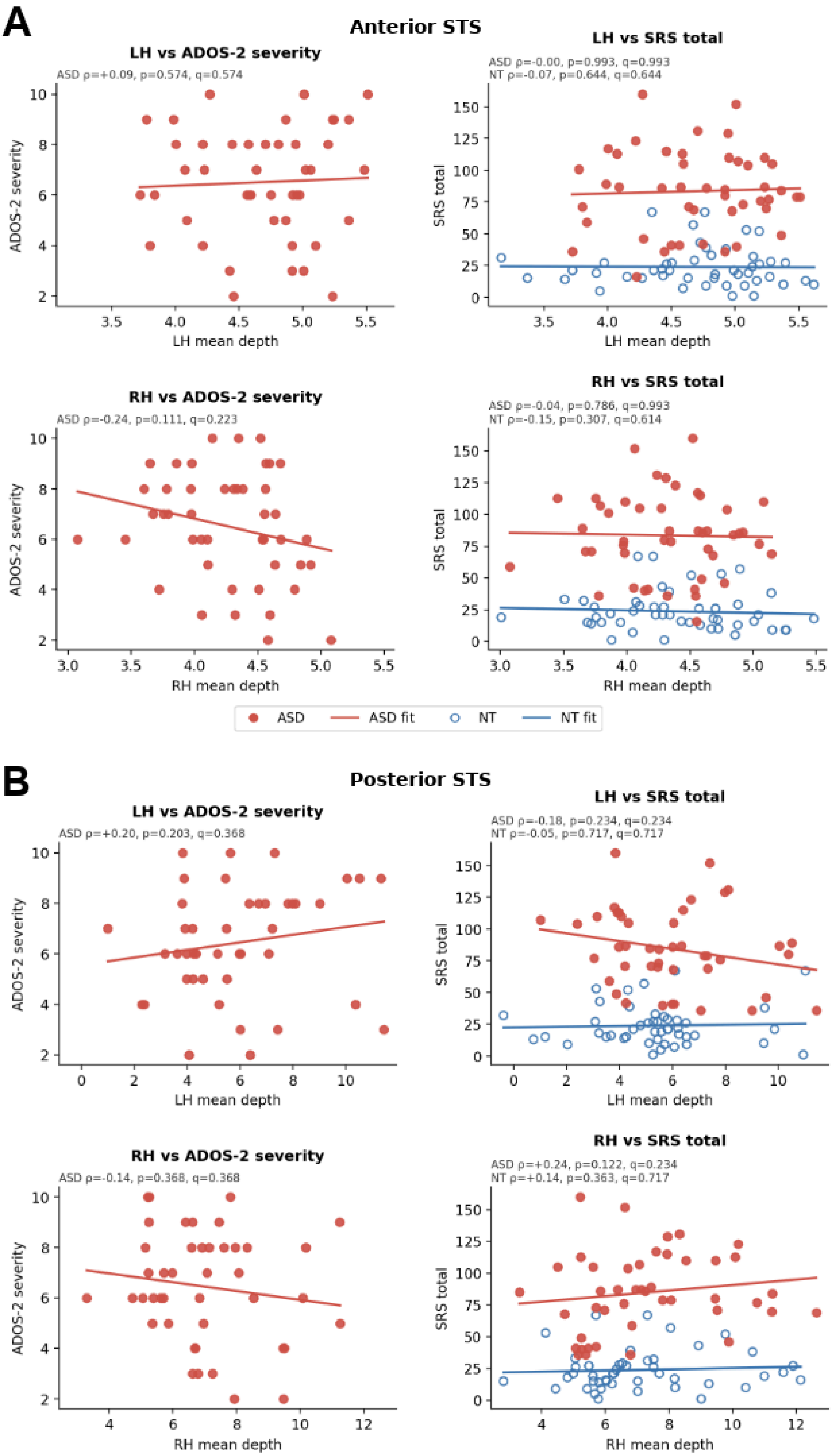
A
Anterior STS
LH vs ADOS-2 severity
ASD ρ=+0.09, p=0.574, q=0.574
ADOS-2 severity
LH mean depth
LH vs SRS total
ASD ρ=-0.00, p=0.993, q=0.993
NT ρ=-0.07, p=0.644, q=0.644
SRS total
LH mean depth
RH vs ADOS-2 severity
ASD ρ=-0.24, p=0.111, q=0.223
ADOS-2 severity
RH mean depth
RH vs SRS total
ASD ρ=-0.04, p=0.786, q=0.993
NT ρ=-0.15, p=0.307, q=0.614
SRS total
RH mean depth
ASD
ASD fit
NT
NT fit
B
Posterior STS
LH vs ADOS-2 severity
ASD ρ=+0.20, p=0.203, q=0.368
ADOS-2 severity
LH mean depth
LH vs SRS total
ASD ρ=-0.18, p=0.234, q=0.234
NT ρ=-0.05, p=0.717, q=0.717
SRS total
LH mean depth
RH vs ADOS-2 severity
ASD ρ=-0.14, p=0.368, q=0.368
ADOS-2 severity
RH mean depth
RH vs SRS total
ASD ρ=+0.24, p=0.122, q=0.234
NT ρ=+0.14, p=0.363, q=0.717
SRS total
RH mean depth

**Figure S6. Mean depth of the anterior and posterior STS does not correlate with ASD severity scores. A.** Scatterplots visualizing the mean depth of the anterior STS (measured in FreeSurfer Units; Dale et al., 1999; Fischl et al., 1999) and its effect on ADOS-2 (left column) and SRS (right column) severity scores. Individual dots represent individual participants (ASD: red, filled; NT: blue, hollow). Solid red (ASD) and blue (NT) lines represent lines of best fit. To control for multiple comparisons across both hemispheres, False Discovery Rate (FDR) correction was applied for each severity score. **B.** Same as **A.**, but for the posterior portion of the STS. LH = left hemisphere, RH = right hemisphere, $\rho$ = Spearman's rank correlation, q = FDR-corrected p-value, Not significant $q > .05$
